# Tissue-specific divergence in sex-biased DNA methylation across the lifespan

**DOI:** 10.64898/2026.04.16.719093

**Authors:** Mandhri D. Abeysooryia, Danielle Hiam, Sarah Voisin, Nir Eynon, Mark Ziemann, Séverine Lamon

## Abstract

**Background:** Ageing is a sex-specific process characterised by a progressive decline in physiological integrity. DNA methylation represents a primary epigenetic hallmark of ageing, yet sex-specific patterns of epigenetic ageing within and across tissues remain poorly understood. This study aims to address these gaps through an integrated analysis of sex-moderated epigenetic ageing across eight human tissues.

**Methods:** A total of 137 DNA methylation datasets comprising over 36,000 individuals aged 10–114 years were analysed using a meta-analytic workflow to identify age-associated differentially methylated positions (aDMPs) and regions (aDMRs), meta-regression to assess sex moderation, and pathway enrichment analyses to interpret functional relevance.

**Findings:** Individual tissues displayed distinct age-related methylation trajectories, but some DMP sites showed consistent hyper- or hypomethylation across tissues. Across tissues, we identified 68,630 aDMPs (10%) robustly associated with ageing. Age-associated changes at the regional level were less common, with only 80 robust age-associated aDMRs detected across tissues, representing 0.09% of analysed regions. Sex moderation was observed for only 16 aDMPs (0.002%), indicating that sex effects on age-associated DNA methylation are largely tissue-specific rather than shared across tissues.

**Interpretation:** Our findings indicate that age-associated DNA methylation changes predominantly occur at isolated CpG sites rather than extended genomic regions and are strongly dependent on tissue and genomic context. The minimal overlap of sex-moderated methylation signals across tissues suggests that age-related sex differences at the epigenetic level are more likely attributable to tissue- and cell-type–specific variation rather than to broadly conserved epigenetic mechanisms shared across tissues.

**Funding:** This study was funded by an Australian Research Council (ARC) Discovery project (DP200101830). Séverine Lamon was funded by an ARC Future Fellowship (FT210100278). Nir Eynon was funded by NHMRC Investigator Grant (APP1194159), and a Hevolution/AFAR New Investigator Award in Aging Biology and Geroscience Research. Mandhri D. Abeysooryia was supported by an Australian Government Research Training Program (RTP) Scholarship.

**Research in context Evidence before this study:** DNA methylation is widely recognised as a central epigenetic hallmark of ageing. Previous research has demonstrated that some age-related methylation changes are conserved across tissues, forming the basis of pan-tissue epigenetic clocks. Most studies to date have primarily examined age effects in isolation. Although biological sex influences ageing trajectories and susceptibility to nearly all age-related diseases, sex-moderated epigenetic ageing has received limited investigation. Specifically, pan-tissue clocks, including GrimAge and PhenoAge, are “sex-aware” but were trained and validated in mixed-sex cohorts, limiting their capacity to disentangle tissue-specific sex effects. Consequently, it remains unclear whether sex-moderated epigenetic ageing signals are shared across tissues or are tissue-specific.

**Added value of this study:** This study provides a large-scale, comprehensive multi-tissue analysis of sex-moderated epigenetic ageing, integrating 137 DNA methylation datasets across eight human tissues and more than 36,000 male and female individuals spanning the lifespan. Our findings show that while age-associated methylation changes are widespread at the CpG level, sex-moderated effects are rare and largely tissue-specific, with minimal overlap across tissues.

**Implications of all the available evidence:** Together, the available evidence indicates that epigenetic ageing is predominantly driven by shared, conserved age-related methylation changes, whereas sex differences in epigenetic ageing are modest and context dependent. These sex-related effects are more likely to reflect tissue- and cell-type–specific variation rather than widespread, shared mechanisms. This underscores the need to develop sex-specific epigenetic clocks and to conduct longitudinal cohort and intervention studies to more precisely characterise sex-specific dynamics of epigenetic ageing across tissues.

## Introduction

Understanding the cellular mechanisms of human ageing is essential for addressing the growing health challenges of an ageing population. Chronological ageing, determined by the amount of time elapsed from an individual’s birth, differs from biological ageing, a more complex process characterized by the progressive decline of cellular, molecular and tissue-specific functions over time. Biological ageing is influenced by a range of factors including sex, health, environment and lifestyle choices, which all shape the epigenome. In particular, sex-specific factors, both biological and environmental, exert significant effects on the biological ageing process (1). Because of its stability and technological accessibility, the DNA methylome has emerged as a key biomarker for estimating the biological age of an individual (2).

Each tissue exhibits unique methylation changes with age (3), but some CpG sites also show consistent hyper- or hypomethylation across tissues as people age (4), providing the basis for pan-tissue epigenetic clocks (5). Sex, on the other hand, has historically been overlooked as a biological variable, despite robust evidence for widespread sex-specific DNA methylation differences that are not confined to X-chromosome inactivation (6).

The original Horvath’s clock does not identify or correct for sex-specific methylation patterns (5), but other widely used pan-tissue clocks, including GrimAge (7, 8) or PhenoAge (9), are “sex-aware”. However, because they were trained and validated in mixed-sex cohorts, these clocks are unable to identify whether specific CpG sites are influenced differently in males versus females, and do not enable identification of tissue-specific sex effects (10). An additional factor limiting our understanding of sex-specific epigenetic changes associated with ageing is that most ageing epigenome-wide association studies (EWAS) treat sex as a confounder rather than as an effect modifier, which can obscure sex-specific signals (11). It follows that sex-specific methylation patterns across ageing remain incompletely characterized despite clear biological relevance (12). Indeed, females generally live longer than males but experience higher morbidity across adulthood, with sex-specific prevalence and presentation of most age-related diseases being the norm rather than the exception (13). Pre-menopausal females also exhibit slower epigenetic ageing compared with age-matched males, although this advantage disappears after menopause (14), suggesting a role of both sex hormones and sex chromosomes in this process (15).

Critical knowledge gaps persist in the characterization of sex-moderated patterns of epigenetic ageing within and across tissues, and sex-specific pan-tissue designs are needed to determine whether sex-moderated ageing signals are shared across tissues or tissue-specific. Yet, apart from a recent GTEx-based analysis reporting contributions of sex and age across nine tissues (16), comprehensive cross-tissue studies of sex-specific epigenetic ageing remain limited.

The study addresses these knowledge gaps through a comprehensive multi-tissue analysis of sex-moderated epigenetic ageing across eight human tissue types. Using meta-analysis, meta-regression and pathway analyses approaches on 137 DNA methylation datasets comprising 36,490 individual samples, we identified 68,630 DNA methylated positions (DMP) robustly associated with ageing across tissues, but only 16 DMPs showing sex moderation ac6ross tissues. This systematic approach reveals fundamental patterns in sex-specific human ageing, offering hypotheses regarding the mechanisms that drive sex differences between and within tissues, and putative opportunities for developing targeted interventions.

## Methods

### Dataset selection and preprocessing

This study was approved by the Deakin University Human Research Ethics Committee (HEAG-H154-2023) and included publicly available, restricted-access, and collaborator-provided DNA methylation datasets identified via systematic searches of PubMed, GEO, and dbGaP. Eligible datasets were generated using Illumina 27K, 450K, or EPIC arrays, included ≥10 healthy participants aged ≥10 years with available age and sex data across skeletal muscle, skin, brain, liver, lung, saliva, blood or adipose tissue, and excluded disease cases to isolate ageing and sex effects. Raw IDAT files underwent standardized quality control, probe filtering, normalization, singular value decomposition, and calculation of β and M values, with full study details documented in a version-controlled GitHub repository (https://github.com/mandhri/Included-studies/blob/master/Included%20studies%20for%20the%20PhD%20project.md).

An outline of the study is presented in Figure 1. Full methods are available in Supplementary Materials.

**Figure 1:**
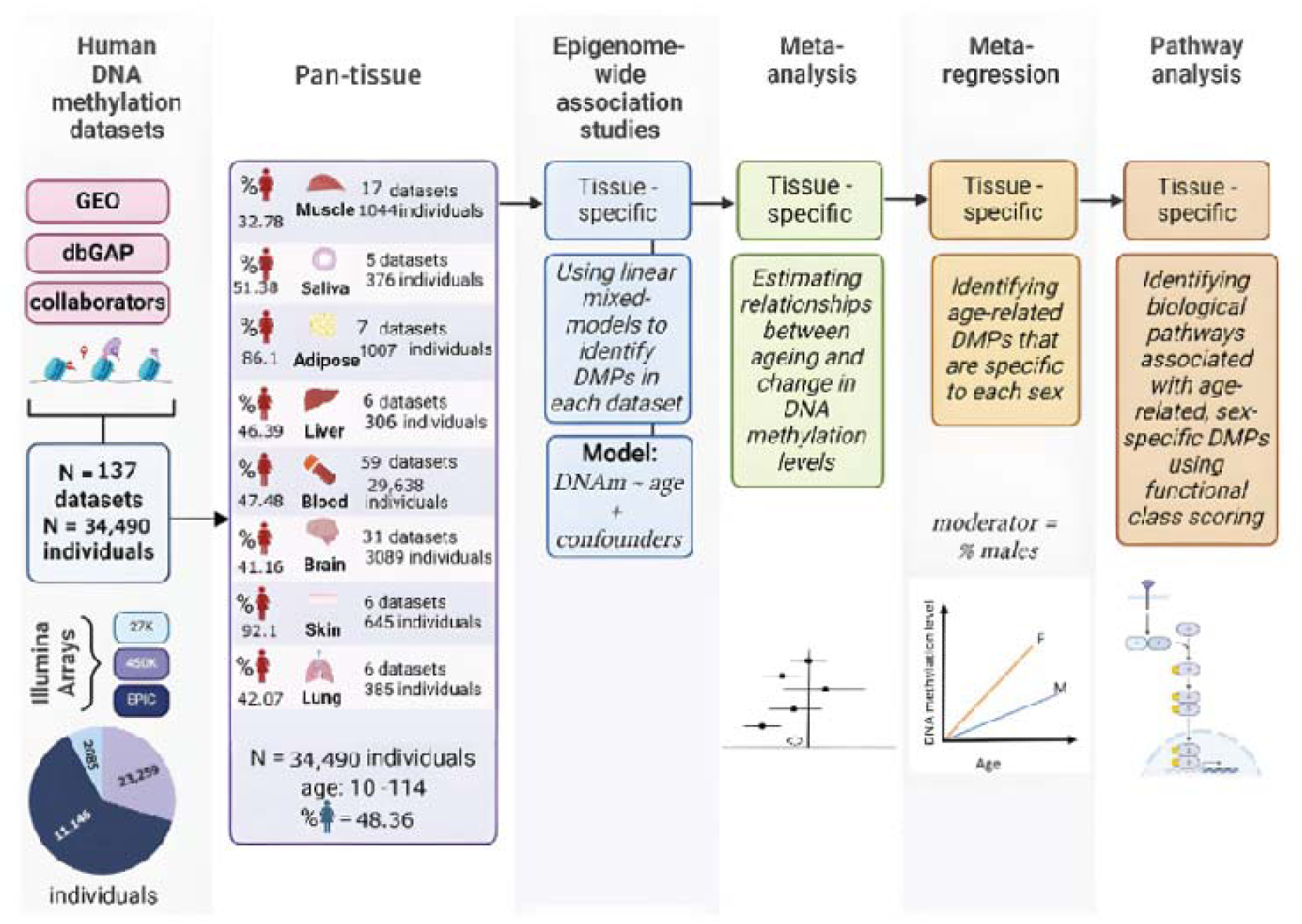
**Schematic overview of the analytical workflow** illustrating the progression from dataset acquisition through epigenome-wide association studies (EWAS), meta-analysis, meta-regression and downstream pathway enrichment for pan-tissue analyses.

### Epigenome-wide association studies (EWAS)

EWAS were conducted using normalized beta values to examine the association between DNA methylation (DNAm) levels and age, with DNAm modelled as the dependent variable and age as the independent variable. Linear models estimated intercept and slope parameters, with the primary aim of identifying age-associated DNAm patterns within each dataset. Covariate adjustment followed a literature-informed approach. Dataset-specific model details are documented in a version-controlled GitHub repository (https://github.com/mandhri/Included-studies/blob/master/ewas_confounders%20PhD%20project.md).

### Random-Effects Meta-Analysis

Age-associated differentially methylated positions (aDMPs) from eight tissue types were pooled for pan-tissue analysis using METAL software (17), which calculated effect sizes and heterogeneity via Cochran’s Q test. A selective filtration criterion was applied wherein aDMPs were retained only if present in ≥30 tissue datasets, ensuring robust cross-tissue patterns. aDMPs were considered significant at FDR < 0.05. Random-effects meta-analysis was performed for each CpG site using the metafor package (18), with empirical Bayes estimation to stabilise variance estimates across studies.

### CpG Selection for Regional Analysis of Differentially Methylated Regions (DMRs)

A regional meta-analysis of DMRs was performed to capture coordinated methylation changes across multiple proximal CpGs. DMR analysis was restricted to CpGs meeting robust representation criteria across cohorts (HetDf ≥ 29 from the METAL output, indicating the presence of a DMR in ≥ 30 tissue datasets). To ensure adequate probe density for regional clustering, DMR analysis was restricted to 450K/EPIC array datasets, excluding 27K array studies due to sparse probe coverage.

### Within-Cohort Correlation Structure and Regional Meta-Analysis

Within each of the eligible 119 cohorts, Pearson correlation matrices of methylation beta values were computed across shared CpGs to account for regional dependence between neighbouring sites. Regional meta-analysis was conducted using the dmrff.meta R package (19), integrating within-cohort correlation structures with random-effects meta-analysis effect sizes and standard errors to identify candidate DMRs, defined as clusters of ≥2 CpGs within 500 bp seeded by CpGs with nominal p < 0.05. DMRs were considered significant at FDR < 0.05, with directionality determined from the meta-analytic effect estimate and supported by z-weighted consistency of constituent CpGs.

### Annotation and Genomic Distribution Analysis

Genomic annotations were derived from TxDb.Hsapiens.UCSC.hg38.knownGene and org.Hs.eg.db, and aDMPs were evaluated across CpG island contexts and functional genomic regions using enrichment analyses. Cross-tissue similarity was quantified using Jaccard indices, and chromosomal distributions were normalized for probe density and chromosome size. Enrichment significance was assessed using Fisher’s exact test with multiple-testing correction.

### Meta-regression

Sex-moderated effects on the DNAm–age association were examined using meta-regression in *metafor*, with the proportion of males per dataset included as a moderator. CpGs showing significant sex moderation (FDR < 0.05) were characterized using the same enrichment framework applied to age-associated aDMPs. For the top signals, linear mixed-effects models incorporating age, sex, and age-by-sex interaction terms (with random intercepts for dataset) were fitted to validate sex-specific methylation trajectories.

### Pathway enrichment analysis

Pathway enrichment analysis was performed in R using the *mitch* package (20, 21), with meta-analysis z-scores as input and gene annotations derived from IlluminaHumanMethylationEPICanno.ilm10b4.hg19, harmonised using HGNChelper.

Gene sets were sourced from MSigDB and Reactome, and pathways were prioritized based on either effect size or statistical significance, retaining those with ≥10 genes.

## Results

### Sample characteristics

A total of 137 DNA methylation datasets comprising 36,490 individual samples from eight tissue types were included in this study. Twelve datasets (10%) had been profiled on Illumina HumanMethylation27 BeadChip arrays (totalling 2085 individuals, 6%), 83 datasets (60%) had been profiled on Illumina HumanMethylation450 BeadChip arrays (totalling 23,259 individuals, 64%) and 42 datasets (30%) had been profiled on Illumina HumanMethylationEPIC BeadChip arrays (totalling 11,146 individuals, 30%).

Participants were aged 10-114 years and 50.9% of them were female. Tissue-specific and sex-specific distribution per tissue are represented in Figure 1.

### Age-associated DMPs (aDMPs) in pan-tissue

EWAS investigating the relationship between DNAm and age (DNAm∼age) were conducted on each of the 137 datasets, through which 874,722 unique CpG sites were detected in at least one dataset. To distinguish between dataset-specific associations and statistically significant age-related probes across all tissues, the datasets were aggregated using METAL and subjected to stringent filtering to determine the minimum number of datasets required to reliably detect robust aDMPs across studies.

A visual examination of the CpG distribution plot produced by METAL identified this inflection point at N = 30 (Supplementary Figure S1). As a result, 664,614 unique CpG sites were identified in 30 or more datasets and selected for further analysis.

### Meta-analysis of aDMPs and aDMRs in pan-tissue

A random-effect meta-analysis was performed on the 664,614 CpGs identified from the EWAS to estimate the association between DNAm and age in pan-tissue. About 10% of CpGs, or 68,630 aDMPs, were differentially methylated with ageing (FDR < 0.05). Of these, 39,169 aDMPs (57%, Figure 2a, red dots) were hypermethylated, and 29,461 aDMPs (43%, Figure 2a, blue dots) were hypomethylated. A histogram of raw p-values distribution is presented in Supplementary Figure S2. The pronounced left-skewed distribution of p-values confirms that the observed associations between age and DNA methylation were unlikely to be due to chance.

**Figure 2a:**
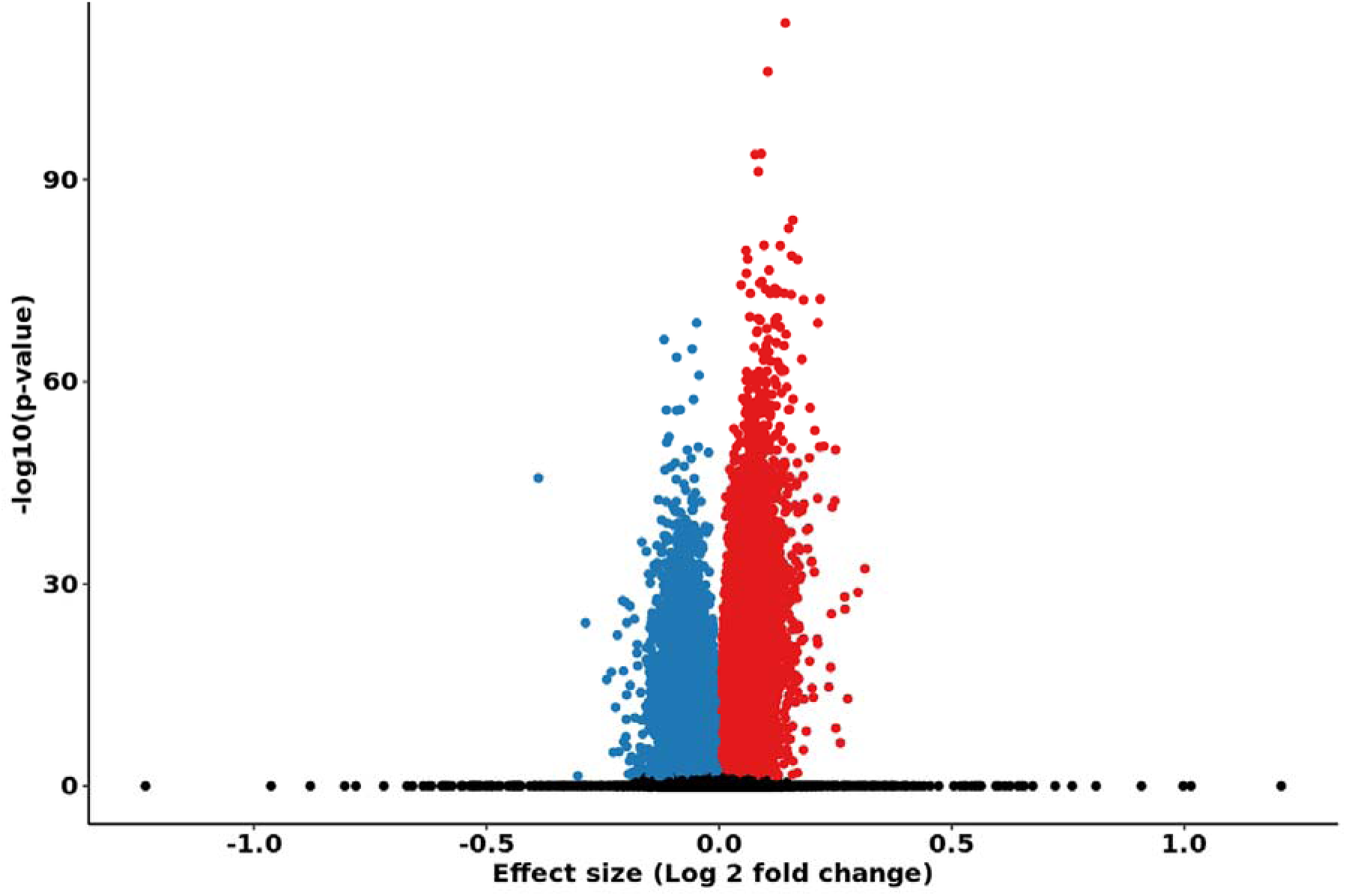
Meta-analysis of age-associated differentially methylated positions (aDMPs) across pan-tissue datasets. The x-axis shows effect size (log fold change), and the y-axis shows statistical significance expressed as −log (adjusted p-value). Red points indicate aDMPs with increased methylation with age, and blue points indicate aDMPs with decreased methylation with age (FDR < 0.05). Black points denote non-significant sites.

Next, a meta-analysis of differentially methylated regions (DMRs) was performed to capture coordinated methylation changes across multiple proximal CpGs, which may have greater biological relevance and improve statistical power compared to single-site analyses. After excluding the studies profiled on Illumina HumanMethylation27 BeadChip due to sparse probe coverage, 8465 unique DMRs were identified in 30 or more datasets and selected for further analysis. Eighty aDMRs (or 0.9%) were differentially methylated with ageing (FDR < 0.05). Of these, 53 aDMRs (66%, Figure 2b, red dots) were hypermethylated, and 27 aDMRs (34%, Figure 2b, blue dots) were hypomethylated.

**Figure 2b:**
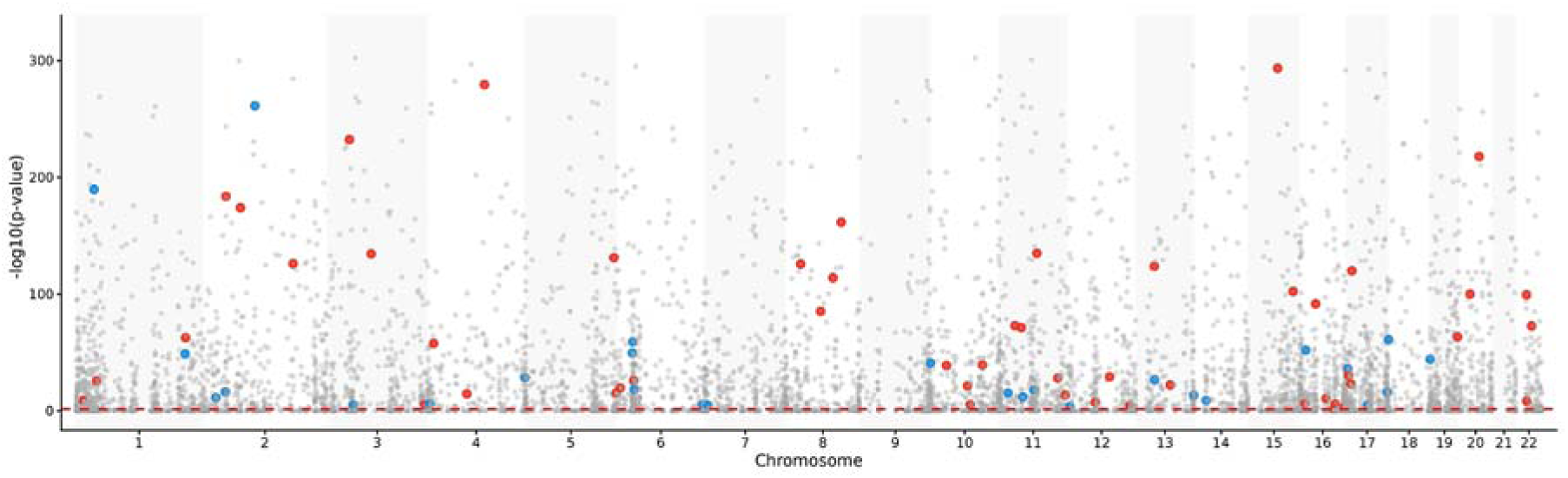
Meta-analysis of age-associated differentially methylated regions (aDMRs) across pan-tissue datasets. The x-axis represents autosomal genomic position, and the y-axis shows statistical significance expressed as −log (adjusted p-value). Red points indicate aDMRs with increased methylation with age, and blue points indicate aDMRs with decreased methylation with age (FDR < 0.05). Grey points denote non-significant sites.

Within the CpG set used for DMR discovery (12,171 CpGs), 2,452 CpGs mapped to significant DMPs (FDR<0.05). Of these, 76 (3.1%) overlapped with a significant DMR and 78 (3.2%) were within ±500 bp of a significant DMR. Conversely, 47/80 significant DMRs (58.8%) contained at least one significant DMP, and all overlapping DMP–DMR pairs showed consistent direction of effect (Supplementary Figure S3).

To determine whether the 68,630 aDMPs identified in pan-tissue reflected changes shared across tissues or were restricted to specific tissues, and to assess the extent to which the pan-tissue aDMP set was recovered within each tissue, parallel meta-analyses were undertaken within each tissue using statistical and methodological parameters identical to the pan-tissue approach. Summary characteristics and results of each individual meta-analyse can be found in Supplementary Table S1.

Jaccard similarity matrices (Figure 3a-c) were then used to visualize the level of similarity between aDMPs in each individual tissue and aDMPs in pan-tissue. Blood tissue demonstrated the highest similarity to pan-tissue, with a Jaccard index of 0.417, indicating that 41.7% of the blood and pan-tissue aDMPs were shared (Figure 3a).

**Figure 3:**
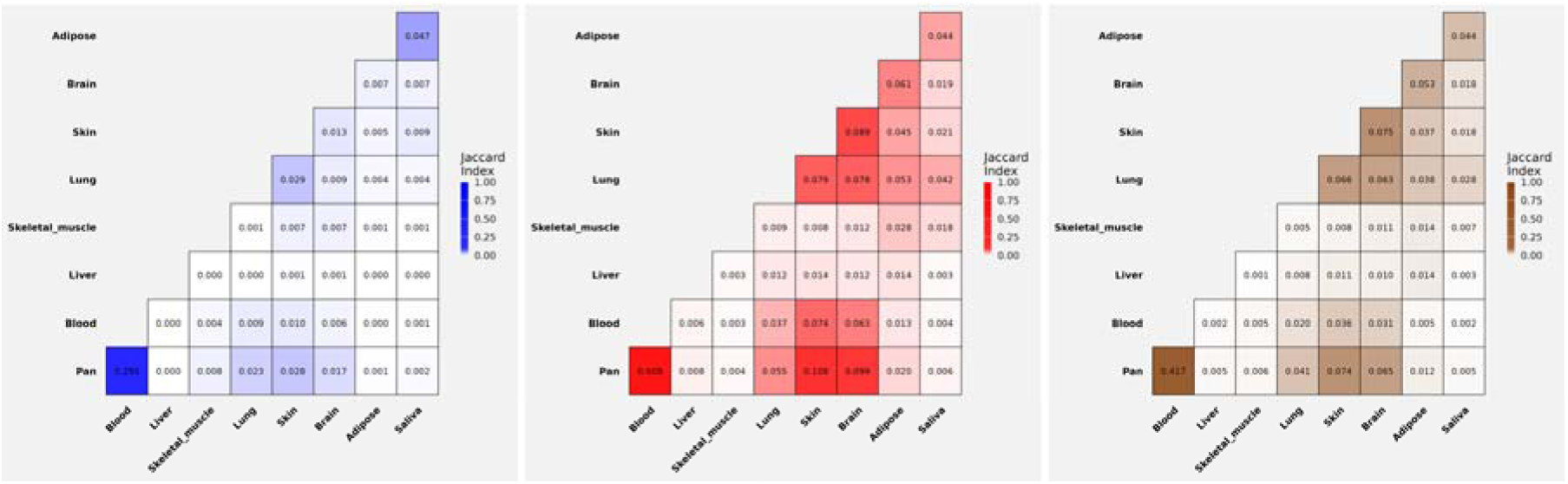
Jaccard similarity matrices showing the overlap of age-associated differentially methylated positions (aDMPs) across tissues. Pairwise Jaccard indices are shown for all significant aDMPs (left panel, blue), hypermethylated aDMPs (middle panel, red), and hypomethylated aDMPs (right panel, brown).

Similar patterns were observed for hyper- and hypomethylated aDMPs (Figure 3b-c).

To further explore the blood’s dominant similarity to the pan-tissue ageing signature, the absolute number of aDMPs (FDR < 0.05) was quantified across each tissue (Supplementary Figure S4). Blood shared 64,430 aDMPs of its 150,356 (42,8%) tissue-specific significant aDMPs with pan-tissue, covering 93.9% (64,430/68,630) of all pan-tissue aDMPs. Other tissues showed substantially lower overlap with pan-tissue despite the detection of many unique aDMPs.

To assess the degree to which the pan-tissue signal was driven by blood tissue, a comparative meta-analysis approach was employed, whereby two separate meta-analyses were performed: one including all tissues, and another in which blood tissue data were excluded. A Pearson correlation of random-effects estimate between the pan-tissue meta-analysis with and without blood yielded an r = 0.71, indicating a moderate-to-strong positive relationship (Supplementary Figure S5).

Excluding blood datasets reduced the number of significant aDMPs with FDR < 0.05 by 85.3% (from 68,630 with blood to 10,091 without blood), confirming that blood tissue substantially amplifies the detection of age-associated methylation sites in the pan-tissue analysis. However, when the overlap between analyses was examined, 88.3% of the 10,091 sites were significant regardless of blood inclusion. This demonstrates that, although blood samples strongly influence the pan-tissue signature, over half of the age-associated methylation effects is maintained when blood data are excluded. Core methylation-age patterns are therefore not entirely blood-dependent.

### Genomic position and chromosomal distribution of aDMPs in pan-tissue

Following the characterization of pan-tissue age-associated methylation patterns, the genomic and functional distribution of aDMPs were quantified to identify the specific regulatory regions targeted by age-related methylation changes (Figure 4a-b, Supplementary Table S2a-b).

**Figure 4.**
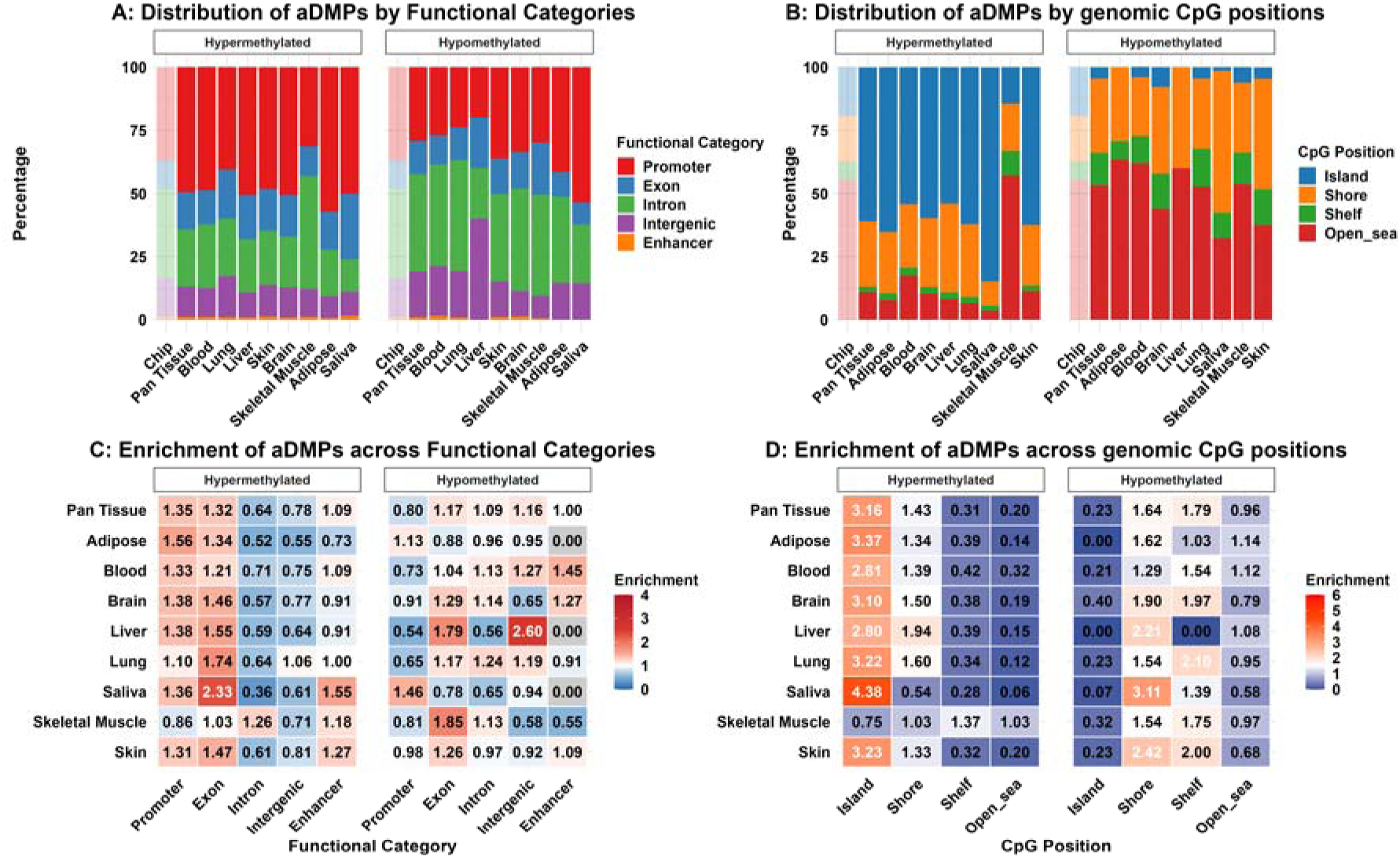
Distribution of age-associated differentially methylated positions (aDMPs) by CpG context and functional genomic annotation across eight tissue types. (A) Proportion of hyper- and hypomethylated aDMPs located in CpG islands, shores, shelves, and open sea regions. (B) Proportion of hyper- and hypomethylated aDMPs located in promoters, exons, introns, intergenic regions, and enhancers. Enrichment analyses are shown relative to the background probe distribution of the array (baseline: islands 19.3%, shores 18.1%, shelves 7.1%, open sea 55.5%; promoters 36.7%, exons 11.2%, introns 35.5%, intergenic regions 15.4%, enhancers 1.1%). Values >1 indicate enrichment (red), values <1 indicate depletion (blue).

The pan-tissue analysis revealed direction-dependent enrichment patterns across CpG contexts. Hypermethylated aDMPs were concentrated in CpG islands, where 60.9% of sites were located despite these regions comprising only 19.3% of array coverage (enrichment ratio: 3.16, FDR ≤ 0.05). Conversely, hypomethylated sites demonstrated clear depletion (4.4%) from CpG islands (enrichment ratio: 0.23, FDR ≤ 0.05) and were more frequent in open sea regions, which contained 53.3% of all hypomethylated aDMPs. This opposing distribution pattern was consistently observed across individual tissues, with blood, brain, and skin most closely matching the pan-tissue hypermethylation profile, while lung and skeletal muscle aligned with the hypomethylation pattern.

A parallel, direction-dependent enrichment was observed in functional genomic annotation, with hypermethylated aDMPs showing a 1.36 enrichment ratio in promoters (46.8% compared with 36.7% background, FDR ≤ 0.05), consistent with their CpG-island concentration. Hypomethylated sites were depleted from promoters (28.7%, FDR ≤ 0.05) and preferentially localised to introns (38.1% compared with 35.5% background, FDR ≤ 0.05) and intergenic regions (17.7% compared with 15.4% background, FDR ≤ 0.05).

Finally, the chromosomal distribution of aDMPs revealed distinct enrichment patterns across the genome (Supplementary Figure S6 A-B). When normalized by probe density, pan-tissue aDMPs demonstrated a relatively balanced distribution of hypermethylation and hypomethylation across most chromosomes, which contrasted with the pronounced hypomethylation bias observed in blood and skeletal muscle and the hypermethylation bias observed in liver, brain, adipose, and skin (Supplementary Figure S6A).

Chromosomes 19 exhibited elevated aDMP densities relative to other chromosomes, paralleling patterns observed in individual tissues. When normalized by chromosomal arm length, substantial enrichment was observed on chromosome 19, with both p and q arms exceeding 300 aDMPs per 10 Mb, representing a 2.8-fold increase over the genome wide average of 107 aDMPs per 10 Mb (Supplementary Fig. S6B). Chromosome 19 enrichment was consistently observed across brain, adipose tissue, lung and skin, indicating a common chromosomal pattern of age-associated methylation changes across tissue types.

### Pathway enrichment analysis

To determine whether the 664,614 CpGs converged on specific biological systems, we performed pathway enrichment analysis using 2,656 Reactome gene sets containing at least 10 genes per set. Of these, 2068 were identified and 231 pathways were significantly enriched at FDR ≤ 0.05. Hypermethylation dominated with 150 hypermethylated pathways (65%) compared to only 81 hypomethylated pathways (35%) (Figure 5).

**Figure 5.**
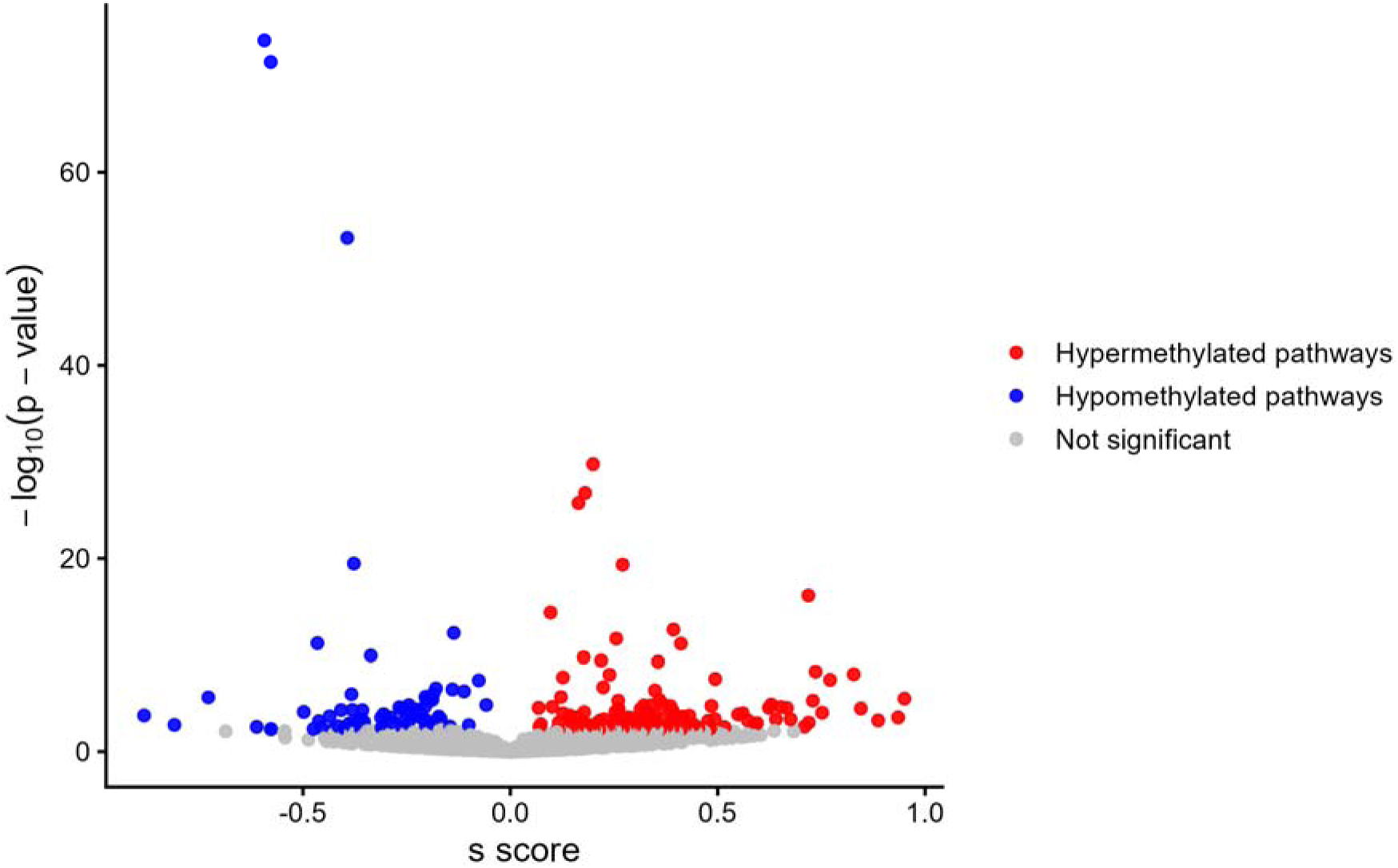
Volcano plot showing gene set enrichment results for age-associated effects across pan-tissue analyses. The x-axis represents the enrichment score (s score), and the y-axis shows statistical significance expressed as −log (p-value). Red points denote significantly enriched hypermethylated gene sets (n = 150; FDR ≤ 0.05), blue points denote significantly enriched hypomethylated gene sets (n = 81; FDR ≤ 0.05), while black points represent non-significant gene sets (n = 1,837).

While at the pathway level, analysis indicated a dominance of hypermethylation, at the gene level, there were more hypomethylated genes (14,338) than hypermethylated genes (7,296) (Supplementary Figure S7). This indicates that hypermethylated genes tend to cluster within specific pathways, whereas hypomethylated genes are dispersed across the genome. The top enriched pathways from the pan-tissue analysis are represented in Supplementary Table S3a and Figure 6.

**Figure 6.**
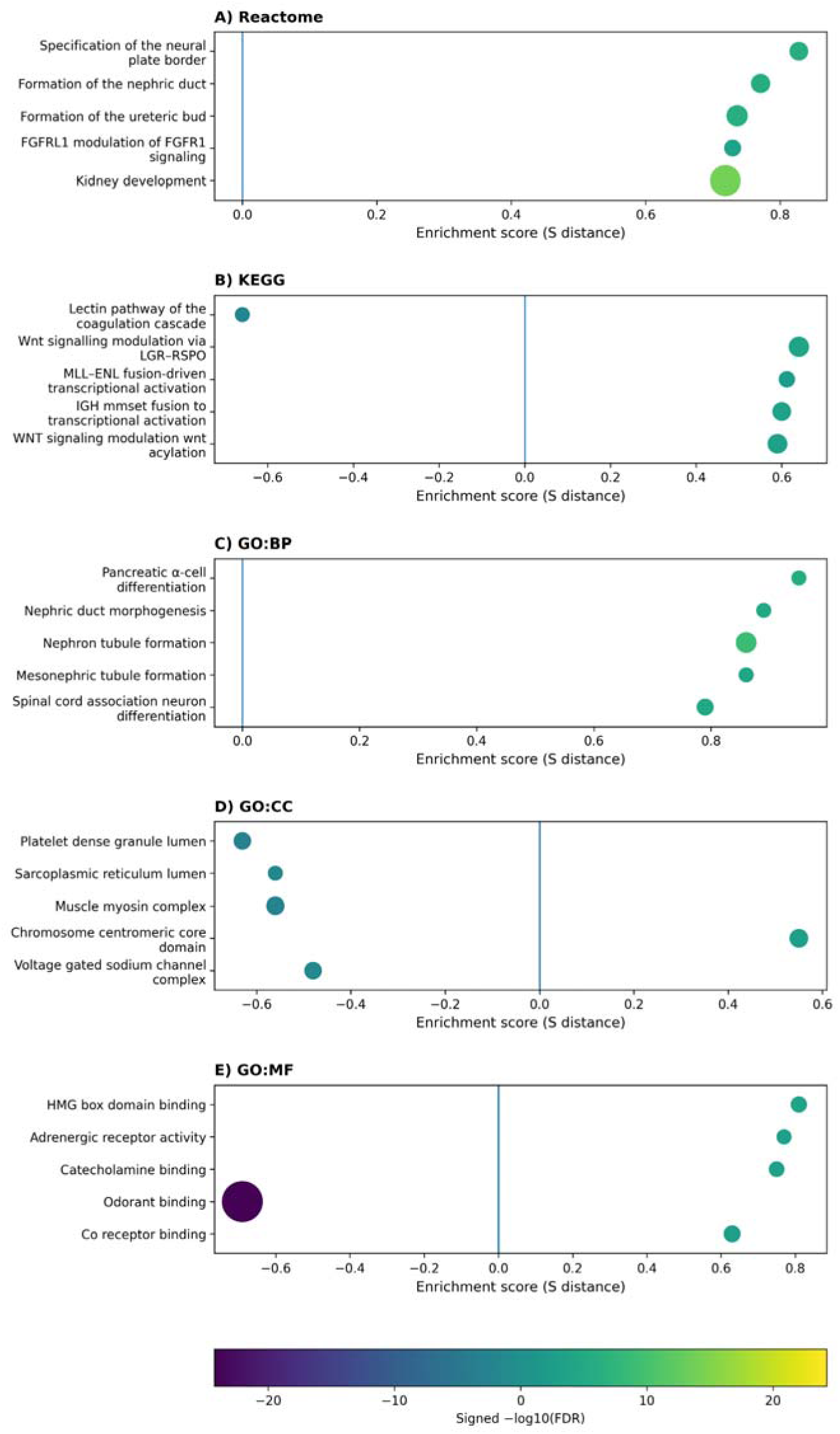
Top-five pathways ranked by smallest absolute enrichment score (s-score) associated with age-related methylation changes across pan-tissue analyses (FDR < 0.05). Pathways are shown for (A) Reactome, (B) KEGG, (C) Gene Ontology (GO) Biological Process (BP), (D) GO Cellular Component (CC), and (E) GO Molecular Function (MF). Only gene sets with a minimum size of n = 10 were included. Dot size reflects gene set size (range: N = 10–84), and colour indicates statistical significance, expressed as signed −log□□(FDR).

Similarly, the 8465 unique DMRs were subjected to pathway enrichment analysis. No pathways were significant for ageing at FDR ≤ 0.05 (Supplementary Table S3b and Supplementary Figure S8).

### Sex moderation of aDMPs in pan-tissue

To determine how sex modifies the associations between sex and DNAm levels in pan-tissue, a meta-regression using the proportion of male in each dataset as a moderator was performed on the 664,614 CpGs selected from the EWAS. Out of 664,614 CpGs, 664,345 returned meta-regression estimates using sex as a moderator prior to FDR correction. A small number of models failed to converge according to the Fisher’s scoring algorithm.

Sixteen sex-moderated aDMPs (0.002%) showed sex moderation across tissues at FDR < 0.05, including five hypermethylated aDMPs (31%) and 11 hypomethylated aDMPs (69%) (Figure 7). A histogram of raw p-values presented in Supplementary Figure S9.

**Figure 7.**
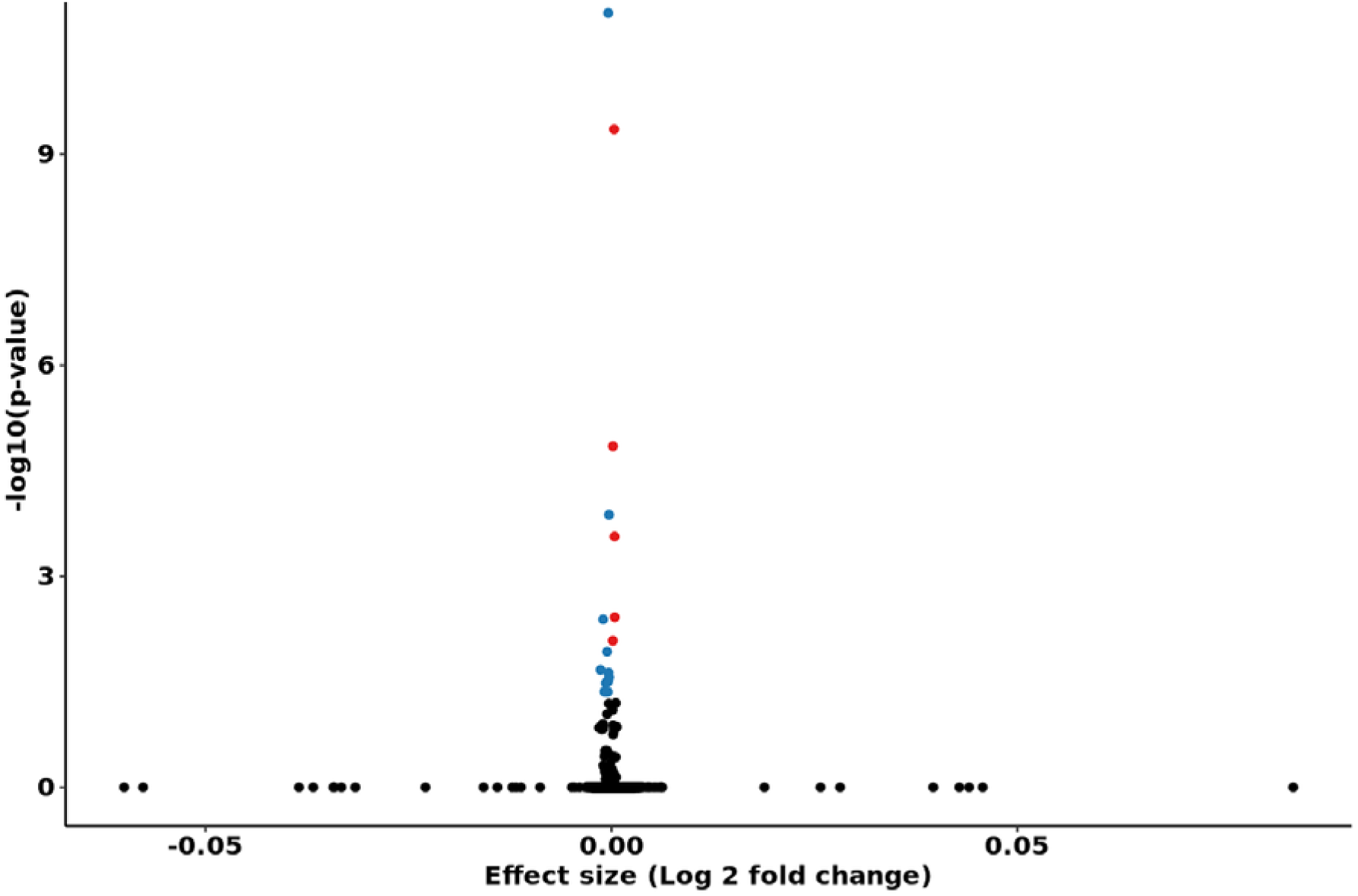
Meta-regression of age-associated differentially methylated positions (aDMPs) across pan-tissue datasets. The x-axis shows effect size (log fold change), and the y-axis shows statistical significance expressed as −log (p-value). Blue points denote aDMPs with decreased methylation associated with the age × sex interaction, red points denote aDMPs with increased methylation, and black points represent non-significant sites (FDR < 0.05).

Given the modest set size (n = 16) of sex associated aDMPs in pan tissue, the genomic and functional context of sex-moderated aDMPs are reported in Supplementary Table S4a-b and Supplementary Figures S10–S12. The Jaccard analysis revealed complete tissue specificity in sex-moderated methylation changes, where the pan-tissue sex-moderated aDMPs showed no overlap with sex-moderated aDMPs from any individual tissue (all Jaccard indices = 0, presented in Supplementary Figure 10). Chromosomal distributions contained few sites per chromosome, reflecting the limited set size (Supplementary Figures S12).

### Interactive modelling of sex-biased aDMPs

Interactive term modelling was applied to the top 10 sex-associated aDMPs identified in the meta-analysis to examine both baseline sex differences and age-by-sex interactions (Figure 8, Supplementary Table S5). Sex-specific methylation trajectories were heterogeneous. Eight of ten CpGs showed a negative age*sex interaction (β <0), indicating that the male methylation trajectory changes more slowly with age than the female trajectory. Theoretical intersection ages ranged from 2 to 151 years, where an early intersection indicates similar DNAm levels at a specific CpG in early life, with trajectories of sex-specific differences displaying different rates thereafter (Supplementary Table S5)

**Figure 8.**
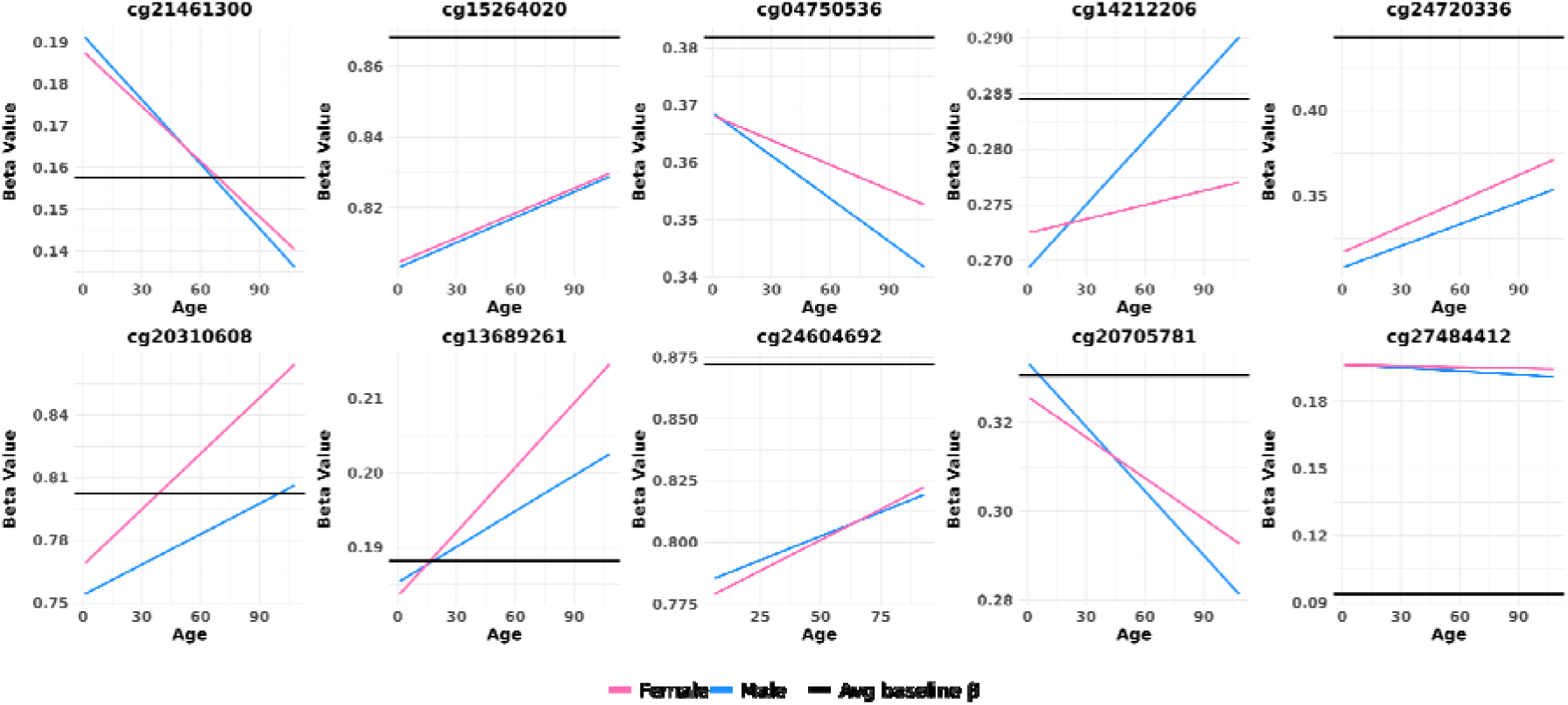
Interactive plots of the top 10 meta-regressed CpG sites identified in pan-tissue analyses at FDR ≤ 0.05. Blue lines show age-related changes in DNA methylation for males, and pink lines show age-related changes for females. Age is shown on the x-axis, and DNA methylation level (β value) is shown on the y-axis. The black line denotes the average β value for each CpG.

## Discussion

This study used a meta-regression approach to analyze 137 DNA methylation datasets from eight human tissues comprising > 36,000 individuals aged 10-114. It identified 68,630 robust age-associated DMPs across tissues, confirming that ageing exerts widespread and reproducible effects in DNA methylation at individual CpG sites across the lifespan. Changes were less consistent at the regional level, with 80 robust age-associated DMRs across tissues. This suggests that most age-associated methylation changes are spatially dispersed or context-dependent, rather than forming large, consistently altered genomic blocks across tissues. Significant DMRs therefore likely represent highly robust and biologically conserved ageing signals.

The ageing signature was heavily weighted by the contribution of the blood methylome, which showed a significant overlap with the pan-tissue signal. This observation likely owes to a combination of two factors: the large representation of blood-derived datasets available in public repositories, and the high cellular turnover of blood tissue (22).

However, sensitivity analysis confirmed a ∼88% overlap among remaining aDMPs after the exclusion of blood datasets, demonstrating the existence of a core, blood-independent ageing methylation programme. This supports the biological validity of pan-tissue clocks, such as Horvath’s clock (5), albeit with some limitations for their interpretation in tissues other than the blood.

Ageing does not affect the methylome randomly instead, it follows direction-conserved rules across tissues. Previous literature shows that age-related hypermethylation predominantly occurs in regulatory regions of the genome (23), while hypomethylation rather occurs in gene-poor and more distal regions (24). Our data support this notion, where hypermethylated aDMP were mostly enriched in CpG islands and promoters, where they may progressively repress developmental genes (23, 25), and hypomethylated aDMP were mostly found in open sea, intronic and intergenic regions. In line with this hypothesis, hypermethylated aDMPs clustered within pathways involved in embryonic development, cell specification and organ differentiation, possibly reflecting a gradual locking-in of differentiated states at the expense of plasticity and regenerative potential. Across tissues, chromosome 19 showed consistently high aDMP density, even after normalisation for probe density and chromosomal length. Chromosome 19 is gene-dense, CpG-rich, and enriched for large families of genes encoding immune, metabolic, and transcriptional regulators (26), which may make it a particularly sensitive candidate to age-related epigenetic remodelling. The predominance of hypermethylated aDMRs mirrors the CpG-level trend and suggests that regional hypermethylation is a defining feature of pan-tissue epigenetic ageing, whereas hypomethylated regions are potentially more tissue- or context-specific.

A surprising finding of this study was that, contrary to our working hypothesis, an extremely small proportion of aDMPs (16, or 0.002%) showed robust moderation by sex across tissues, suggesting a near-complete tissue specificity of sex effects. This result demonstrates that differences in epigenetic ageing are therefore driven by tissue-specific effects over constitutive variability at the CpG level. This might also reflect higher level regulatory mechanisms that cannot captured at the individual CpG resolution, including alterations of chromatin state or enhancer network dynamics. Within the top 10 sex-moderated CpGs, most sites displayed slower age-related changes in methylation in males than in females despite heterogenous baseline levels. These differences might reflect the influence of sex hormone variations across the lifespan, but also the sex-specific accumulation of exposure to lifestyle (27) and environmental (28) factors. The wide range of theoretical intersection ages (2–151 years) further highlights that, when present, sex differences in methylation accumulate gradually, rather than being fixed early in life.

The minimal overlap in sex moderated aDMPS was unexpected, as it contrasts with well-established sex differences in lifespan and healthspan (29), disease risk and prevalence (13), tissue-specific ageing processes (30) and epigenetic clock acceleration (31). Our results suggest that, at the epigenetic level, age-related sex differences may therefore be rather driven by cell-type composition, which represents a known limitation of the analysis of bulk tissue methylation datasets when no cell type correction or sensitivity analysis can be performed (32). This is particularly relevant for tissues including blood, skeletal muscle, brain and adipose tissue, where cellular composition changes substantially with age and differs between sexes (1). However, other epigenetic mechanisms, including non-coding RNA regulation and histone accessibility may also be major players underlying lifelong sex differences in epigenetic ageing. Overall, sex differences in biological ageing might be driven by small, cumulative epigenetic effects, which may only become biologically meaningful at the systems or tissue-function level.

A limitation of the current study is that CpG sites located on the X and Y chromosomes were excluded during pre-processing. The X chromosome harbours strong sex-associated methylation differences due to X-chromosome inactivation and dosage effects. Currently, we are unaware of any analytical approach that can capture potential sex-moderated ageing patterns occurring at X-linked loci. However, restricting analyses to autosomal CpGs also provides a conservative test of sex moderation that is not driven by known large-scale methylation differences arising from X-chromosome inactivation. The observation that only a small number of autosomal aDMPs demonstrated sex moderation across tissues is more likely attributable to tissue- and cell-type–specific variation rather than to broadly conserved, sex-specific epigenetic mechanisms shared across tissues.

In conclusion, ageing exerts strong, reproducible effects on the human methylome across tissues, but sex effects in ageing are context-, tissue- and mechanism-dependant. Treating sex as a global modulator of epigenetic ageing may be overly simplistic, as sex aDMPs seem to vary across tissues. This warrants the development of more sex-aware epigenetic clocks such as the sex-adjusted GrimAge/GrimAge2 (7, 8) and PhenoAge (9) clocks.

Finally, while the scale and tissue diversity of this study demonstrate the power of meta-analytic approaches in pan-tissue, the cross-sectional nature of most included datasets limits causal inference. Longitudinal cohort and intervention studies would be required to more precisely characterise sex-specific dynamics of epigenetic ageing across human tissues.

## Ethics approval

This study received ethics approval from the Deakin University’s Human Ethics Committee (HEAG-H154-2023).

## Contributors

MDA curated the data, completed the statistical analysis, interpreted the data and drafted the manuscript. DSA curated the data and completed the statistical analysis. SV and NE conceived/designed the study. MZ completed the statistical analysis and interpreted the data. SL conceived/designed the study, interpreted the data and drafted the manuscript. SL and NE funded the study. All authors critically revised the manuscript, approved the final version, and agreed to be accountable for all aspects of the work.

## Data sharing

The list of datasets used in this analysis is available at Included-studies/Included studies for the PhD project.md at master · mandhri/Included-studies · GitHub

## Acknowledgments

This study was funded by an Australian Research Council (ARC) Discovery project (DP200101830). Séverine Lamon was funded by an ARC Future Fellowship (FT210100278). Nir Eynon was funded by NHMRC Investigator Grant (APP1194159), and a Hevolution/AFAR New Investigator Award in Aging Biology and Geroscience Research. Mandhri D. Abeysooryia was supported by an Australian Government Research Training Program (RTP) Scholarship.

## Supplementary Methods

### Dataset selection and preprocessing

This study received ethics approval from the Deakin University’s Human Ethics Committee (HEAG-H154-2023). Publicly available and restricted access DNA methylation datasets were used in this study. Datasets were selected from a PubMed search conducted using keywords including “Infinium” AND “methylation”; “Pan-tissue” AND “pan-tissue”; “Tissue” AND “methylation” AND “age”; “methylation” AND “tissue” AND “sex and retrieved from GEO or dbGAP. Additional, unpublished datasets were obtained directly from collaborators. Inclusion criteria included datasets profiled on Illumina HumanMethylation27 BeadChip arrays, Illumina HumanMethylation450 BeadChip arrays, or Illumina HumanMethylationEPIC BeadChip arrays (Illumina Inc., San Diego, USA) and containing at least 10 healthy human participants of any sex for which one or several human tissues were analyzed, including skeletal muscle, skin, brain, liver, lung, saliva, blood or adipose tissue; datasets where age, sex and health status (disease) information was available. Participants < 10 years of age were excluded. To focus on the direct effects of sex on DNA methylation during ageing without the interference of disease, only healthy participants (including healthy control participants in the case of disease studies) were selected for analysis. During preprocessing, probes mapping to the X and Y chromosomes were removed and analyses were restricted to autosomal CpG sites. This approach follows common preprocessing practices in epigenome-wide association studies to avoid confounding arising from sex-chromosome dosage differences and X-chromosome inactivation patterns.

A full list of selected studies and related information is recorded in a version-controlled GitHub repository: https://github.com/mandhri/Included-studies/blob/master/Included%20studies%20for%20the%20PhD%20project.md.

The raw methylation data were downloaded in the form of IDAT files and underwent a comprehensive quality control process, including filtration of probes, normalization, singular value decomposition, and calculation of β and M values.

#### Quality control

Low-quality samples were excluded if more than 10% of probes failed detection (defined as a detection p-value > 0.01). Probes were removed if they had missing beta values (detection p-value > 0.01) or a bead count < 3 in at least 5% of samples within a given dataset. Probes targeting non-CpG sites or mapping to multiple genomic locations were also excluded.

#### Exclusion of sex chromosome probes

Probes mapping to the sex chromosomes (X and Y) were excluded due to unequal copy number between males and females, X-chromosome inactivation, and comparatively heterogeneous probe coverage. These factors may introduce methylation differences unrelated to age. Excluding sex chromosome probes therefore minimises confounding when comparing age-associated autosomal methylation patterns between sexes.

#### Exclusion of probes overlapping known single-nucleotide polymorphisms (SNPs)

Probes overlapping known SNPs were filtered using the champ.filter function, based on the built-in SNP annotation lists provided in ChAMP (version 2.32.0) for each array type (27k, 450k, and EPIC) (33). These annotation lists are derived from (34), which defines SNPs as variants catalogued in the 1000 Genomes Project with a global minor allele frequency (MAF) > 1% and located within five bases of the CpG interrogation or single-base extension site. By default, the champ.filter function removes probes overlapping such variants, thereby reducing the likelihood of confounding by common genetic variation. This step ensures that downstream analyses reflect true epigenetic variation rather than underlying genetic differences.

To further enhance data specificity, probes with the potential to hybridise to multiple genomic locations (cross-hybridising probes) were removed using a customised list from(35), as the list included in the current version of ChAMP is outdated.

### Epigenome-wide association studies (EWAS)

The normalised beta values and related phenotypic files of each dataset were used to test the association between DNAm level and age. In the EWAS model, DNAm level is the response (dependent) variable, whereas age is the explanatory (independent) variable.

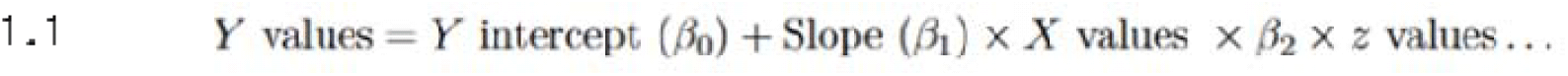

The general statistical EWAS model equation used in the analysis was:

In equation 1.1, β_0_ and β_1_were the EWAS model parameters that were estimated during the modelling. β_0_ represents the Y intercept of the graph, which illustrated the potential value of the dependent variable (DNAm) when the independent variable (age) is zero. β_1_ represents the slope of the curve, which reflects the magnitude of the change in the dependent variable per one unit of change in the independent variable.

The main goal of the EWAS model of this study was to observe how differences in age affects individual DNAm patterns in a given dataset. The collected datasets exhibited heterogeneity in terms of measured variables, where some datasets contained relevant variables other than age that may be associated with DNAm levels (e.g., sex, BMI, alcohol consumption, smoking status, hormone replacement therapy etc) according to the literature, while others included twin studies or repeated measures from clinical studies. Covariate adjustment followed a literature-driven approach, where only variables with established evidence of association with the DNAm relationship were incorporated into the model. Information on the EWAS equation used for each dataset, sample size after filtering for disease (sample size), name of the disease (if present in the dataset along with controls) and tissue/cell-type is recorded in a version-controlled GitHub repository: https://github.com/mandhri/Included-studies/blob/master/ewas_confounders%20PhD%20project.md.

### Within-Cohort Correlation Structure and Regional Meta-Analysis

For each of the 119 cohorts, within-cohort Pearson correlation matrices of methylation beta values were computed across the common CpG sites. Such matrices capture the linkage disequilibrium structure of DNA methylation and are essential for correcting regional test statistics for regional dependence between nearby CpGs.

Regional meta-analysis was performed using the R package dmrff.meta (19), which implements correlation-adjusted regional hypothesis testing. For each cohort, dmrff.meta used the within-cohort correlation matrices combined with the random-effects meta-analysis effect size (ES) and standard errors (SE) from the metafor analysis to identify candidate differentially methylated regions (DMRs).

Candidate DMRs were defined as clusters of CpGs separated by ≤500 bp, seeded by CpGs with nominal p-value < 0.05 in the random-effects meta-analysis. Regional test statistics were calculated using correlation-adjusted inverse-variance weighting, accounting for the covariance structure of methylation changes across the clustered CpGs. DMRs were considered statistically significant at FDR < 0.05 (Bonferroni correction) and required ≥ 2 constituent CpGs per region. The direction of methylation changes for each was determined from the meta-analysis effect estimate. Positive effect estimates (ES > 0) indicated hypermethylation with increasing age, whereas negative estimates (ES < 0) indicated hypomethylation with increasing age.

To assess whether CpGs within each significant DMR supported a consistent direction of age-associated methylation change, DMRs (defined as ≥2 CpGs, FDR<0.05) were mapped to their constituent CpGs using dmrff.sites() and CpG coordinates from the dmrff EWAS output. CpG direction was defined by the sign of the CpG-level effect estimate, and DMR direction by the sign of the DMR estimate. For each DMR, a z-weighted direction score was calculated using the proportion of CpGs whose direction matched the DMR direction, giving more weight to CpGs with larger absolute z-scores (z) so that CpGs with stronger evidence contributed more to the score.

### Annotation and Genomic Distribution Analysis

Genomic annotations were obtained from TxDb.Hsapiens.UCSC.hg38.knownGene and org.Hs.eg.db. The distribution of aDMPs was analysed across CpG island contexts (islands, shores, shelves, open seas) and functional regions (promoters, exons, introns, enhancers, intergenic regions). For each context, enrichment values were calculated using the following formula:

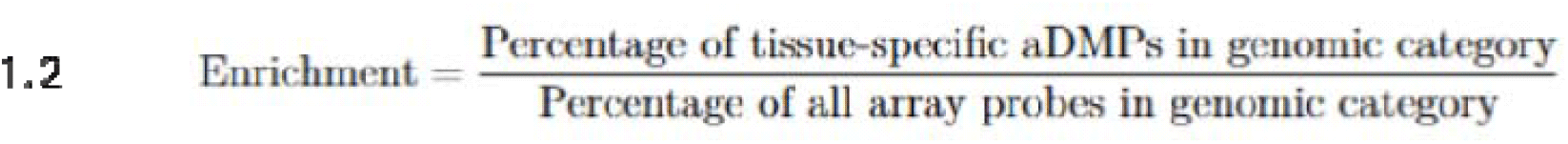

Cross-tissue similarity was assessed using Jaccard indices, calculated separately for hypermethylated and hypomethylated sites.

Chromosomal distribution analysis incorporated normalisation for both probe density and chromosome size. aDMP density was calculated as:

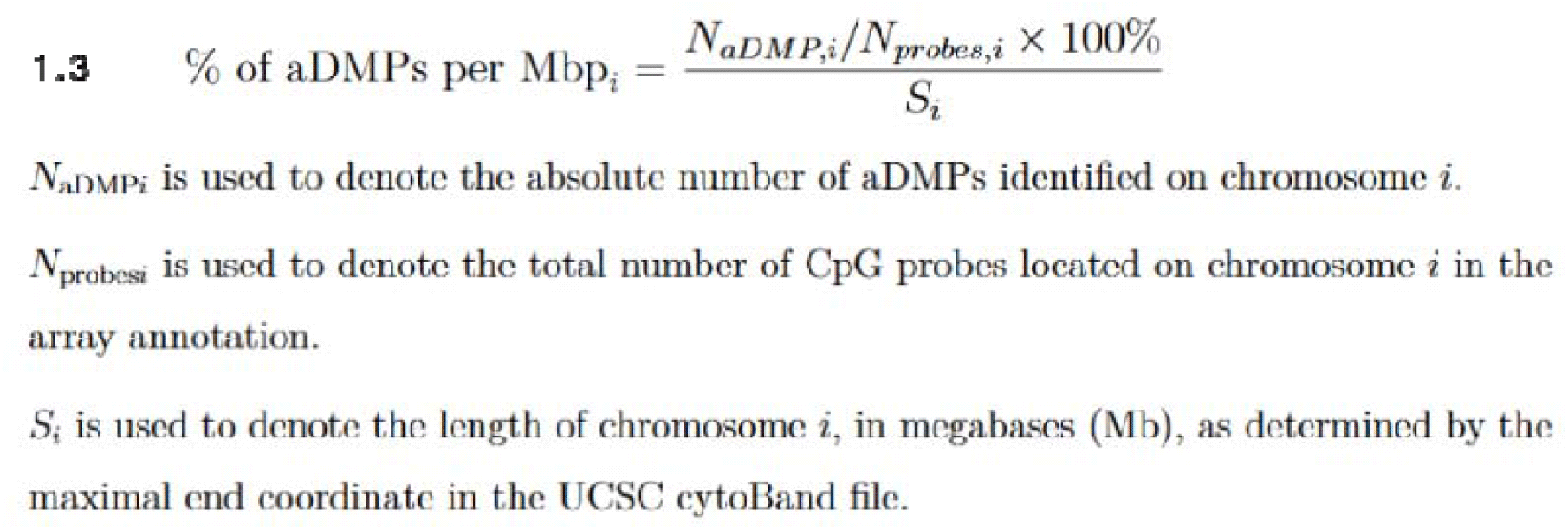

Enrichment ratios were assessed using Fisher’s exact test and resulting p-values were adjusted for multiple comparisons.

## Meta-regression

Sex-moderated effects on the DNAm∼age relationship were examined through meta-regression using the proportion of males per dataset as the moderator variable. Mixed linear models were fitted using the metafor package.

The distribution and enrichment patterns of sex-moderated aDMPs followed the same analytical framework as age-associated aDMPs, with analyses focused on CpGs exhibiting significant sex-moderation (FDR < 0.05).

Interactive modelling validated sex-specific methylation trajectories for the top 10 sex-moderated aDMPs. Linear mixed-effects models were fitted incorporating age, sex, and age-by-sex interaction terms, with random intercepts for datasets to account for between-study variability:

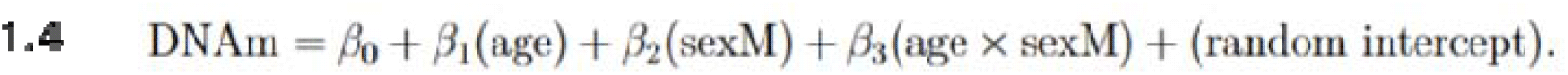

β1 and β0 capture the intercept and slope for females, respectively, whereas β2 and β3 capture how males differ from females in intercept and slope, respectively. The interaction term enabled identification of potential crossing points where male and female methylation trajectories intersect.

### Pathway enrichment analysis

Pathway enrichment analysis was performed using the mitch package (20, 21) in R to identify differentially enriched pathways in pan tissue. Gene annotations were sourced from Gene annotations were sourced from the IlluminaHumanMethylationEPICanno.ilm10b4.hg19 annotation package (36), which provides detailed annotations for the Illumina Human Methylation EPIC array, mapping CpG probes to gene symbols. To standardise gene symbols and correct outdated annotations, the *HGNChelper* package (version 0.8.14) (37) was used, which frequently updates gene symbols according to the latest HUGO Gene Nomenclature Committee (HGNC) guidelines. Z-scores calculated from the meta-analysis results served as input statistics for pathway enrichment. Two prioritising strategies were employed: Effect-size Prioritisation where prioritised pathways were based on the magnitude of the enrichment effect (priority=“effect”), and statistical significance prioritisation where prioritised pathways were based on the statistical significance of the enrichment *(priority = “significance”).* For each strategy, only pathways containing at least five genes (*minsetsize = 5), FDR < 0.05 and absolute standardised enrichment scores(ES) > 0.3* were included to enhance statistical robustness. Duplicated pathways resulting from the two prioritisation strategies were identified and consolidated by retaining unique entries based on pathway name, size, p-values, and enrichment scores. The gene sets required for pathway enrichment analysis were obtained from the Molecular Signatures Database (MSigDB) via the *msigdb* R package, including Gene Ontology Biological Process (GO:BP), Cellular Component (GO:CC), Molecular Function (GO:MF), and KEGG pathway collections. Reactome pathway gene sets were obtained directly from the Reactome pathways database (https://academic.oup.com/nar/article/52/D1/D672/7369850, downloaded on 11 January 2024).

## Supplementary Figures

**Supplementary Figure S1.**
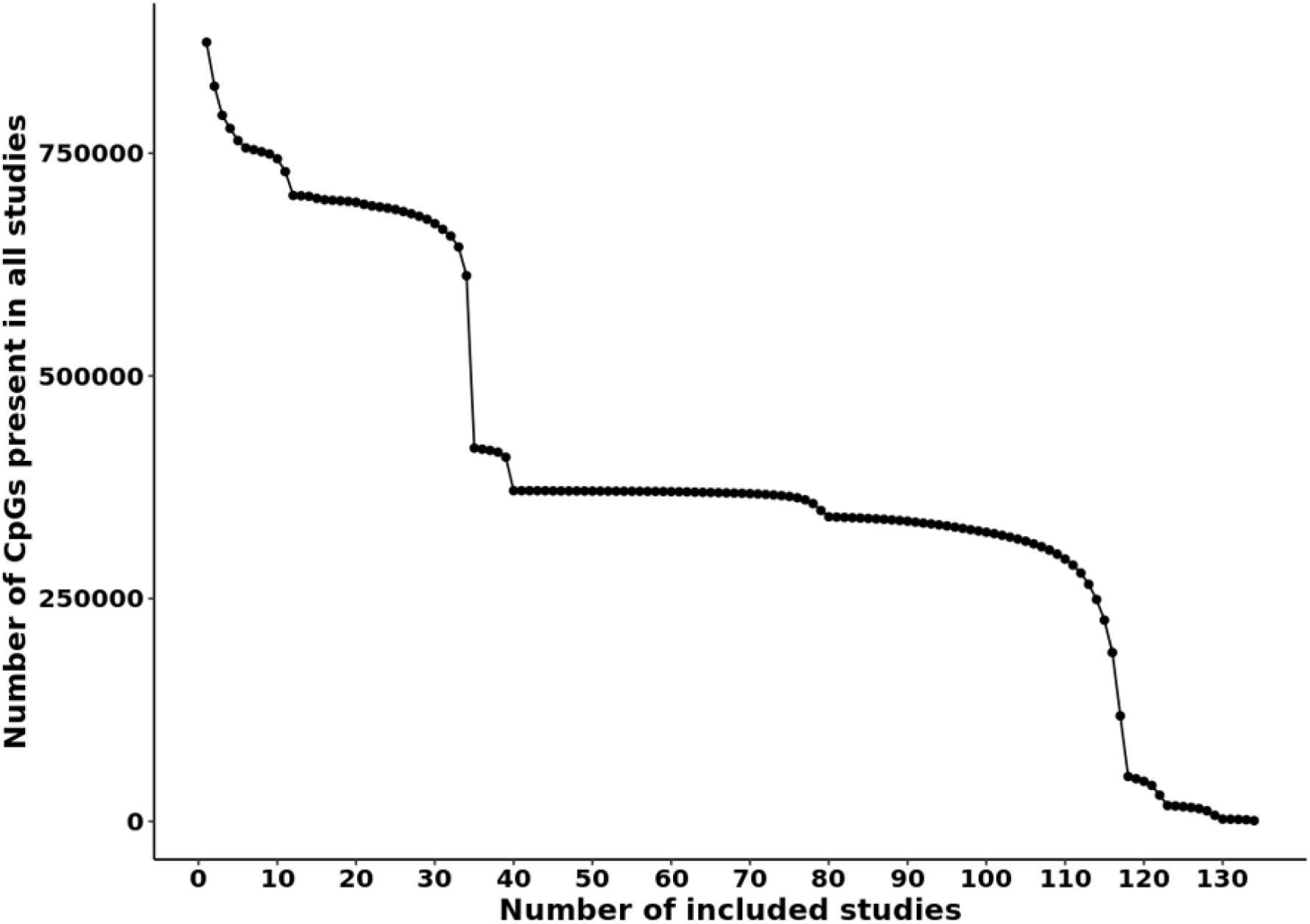
Minimum number of datasets required for robust pan-tissue aDMP detection. Cumulative number of CpG sites retained as a function of the number of included tissue datasets in the METAL meta-analysis. The inflection point at *N* = 30 datasets (vertical drop) was used as a data-driven threshold to define robustly represented CpGs for downstream pan-tissue analyses.

**Supplementary Figure S2.**
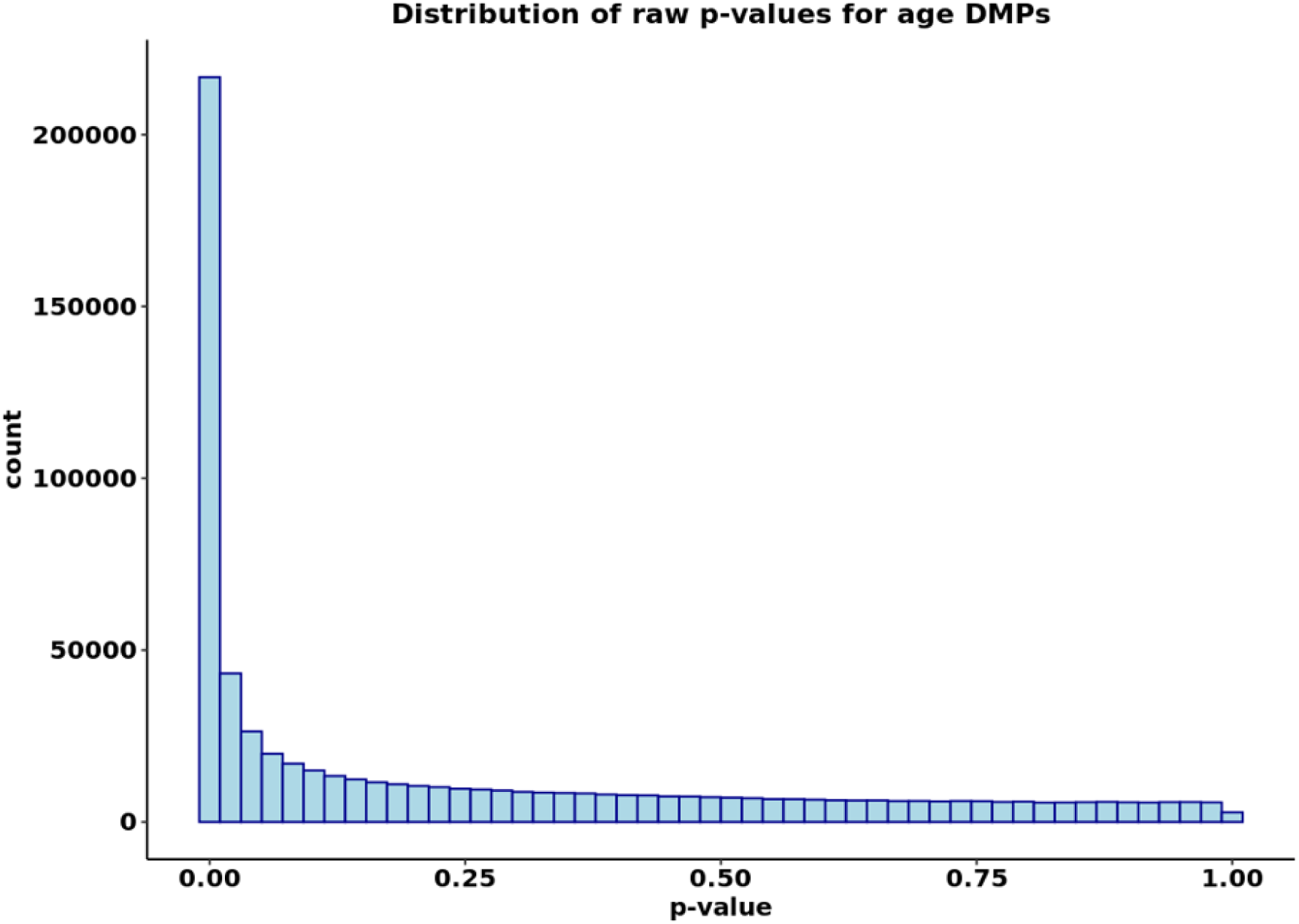
Distribution of raw p-values for age-associated differentially methylated positions (aDMPs). Histogram of unadjusted p-values from the random-effects meta-analysis of DNA methylation ∼ age across pan-tissue datasets. The pronounced left-skew indicates widespread age-associated methylation changes beyond that expected under the null hypothesis.

**Supplementary Figure S3.**
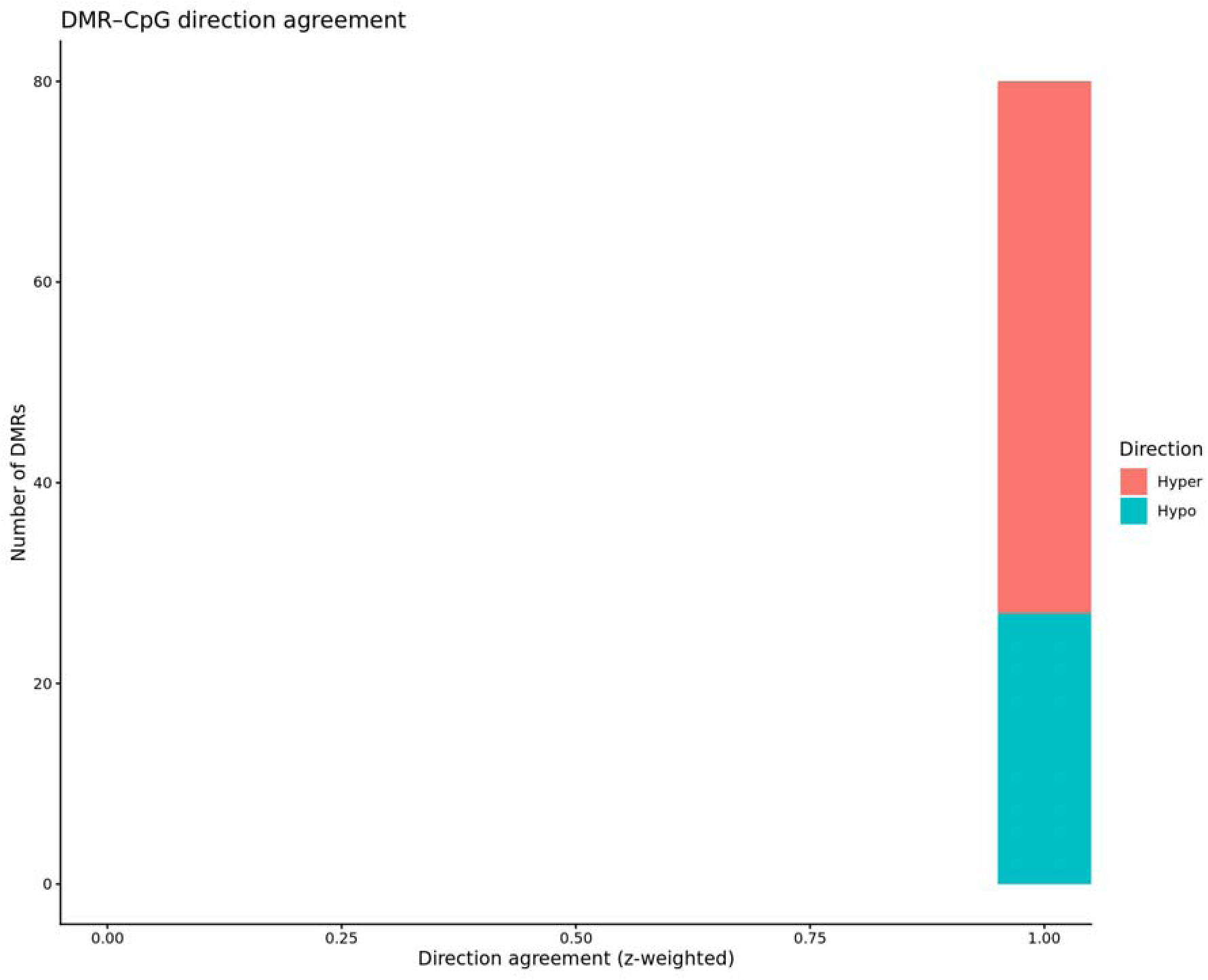
DMR – CpG direction agreement. Red indicates hypermethylation. Green indicates hypomethylation.

**Supplementary Figure S4.**
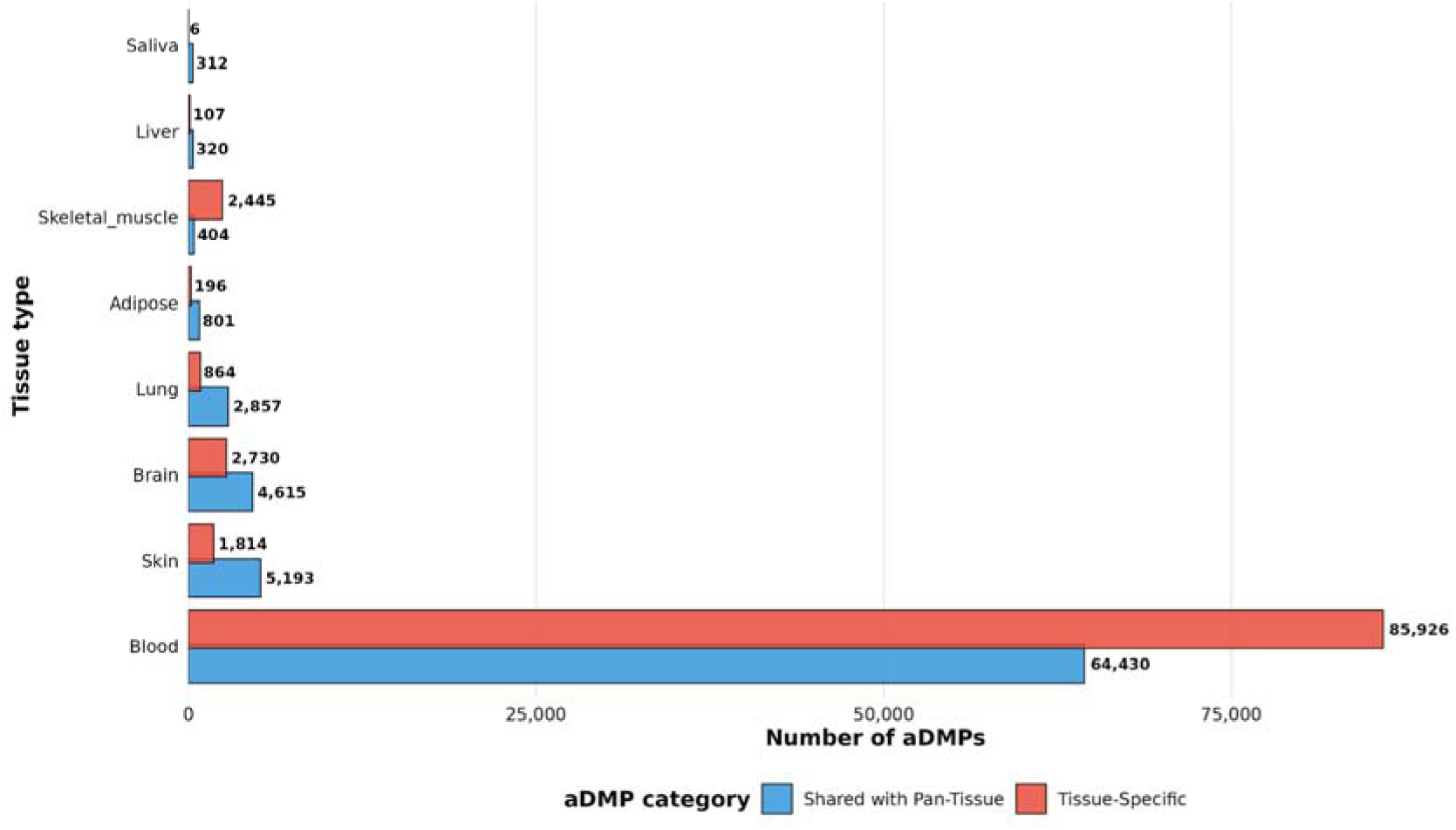
Absolute number of significant aDMPs detected per tissue. Bar plot showing the number of age-associated differentially methylated positions (FDR < 0.05) identified in each tissue-specific meta-analysis, alongside the overlap with pan-tissue aDMPs. Blood demonstrates the highest recovery of pan-tissue aDMPs.

**Supplementary Figure S5.**
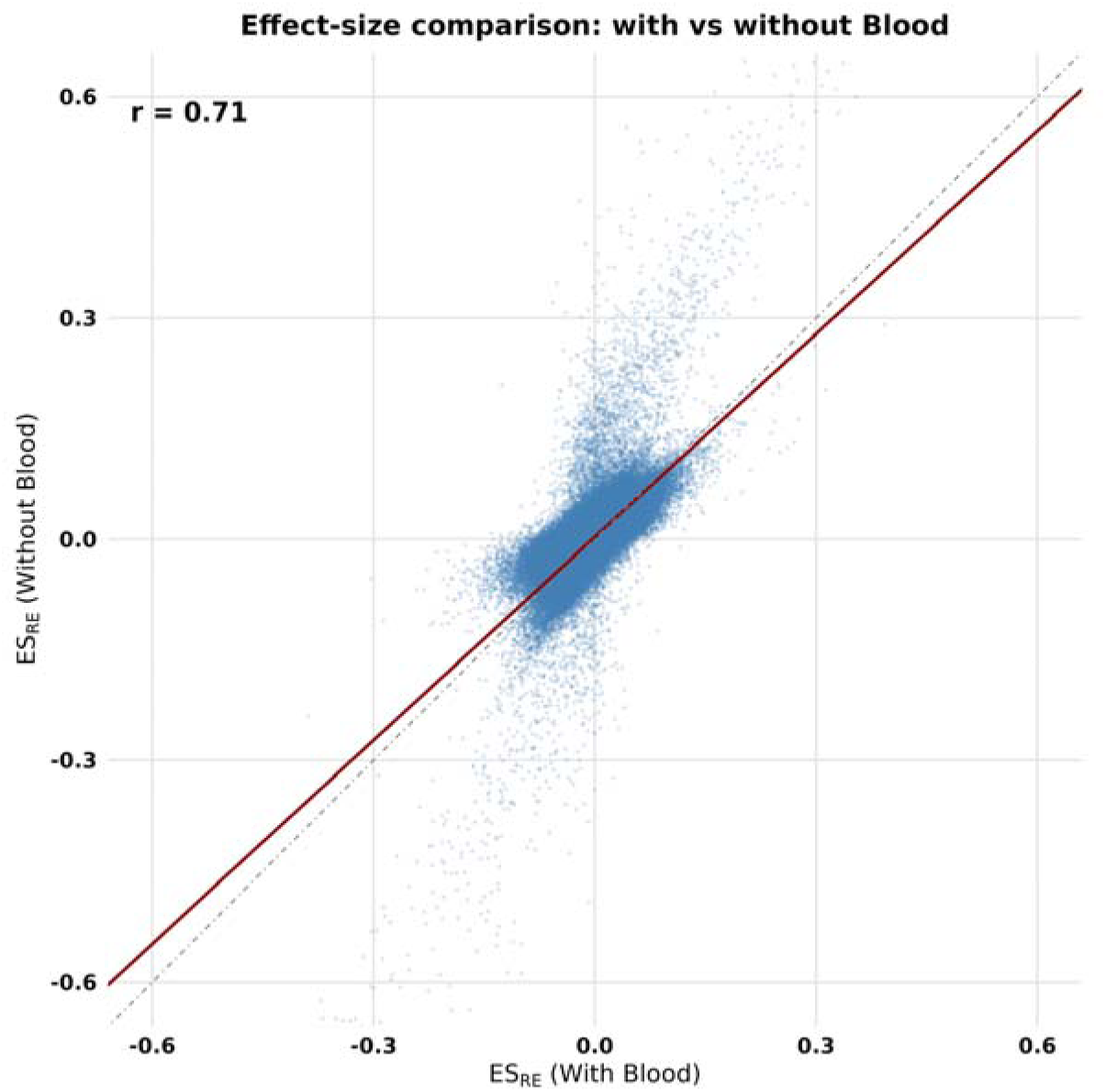
Effect size comparison of pan-tissue aDMPs with and without blood inclusion. Scatter plot comparing random-effects effect size estimates (ES_RE) from pan-tissue meta-analyses performed with all tissues included versus excluding blood datasets. The dashed line represents identity (y = x). Pearson correlation coefficient (*r* = 0.71) indicates moderate-to-strong concordance between models.

**Supplementary Figure S6.**
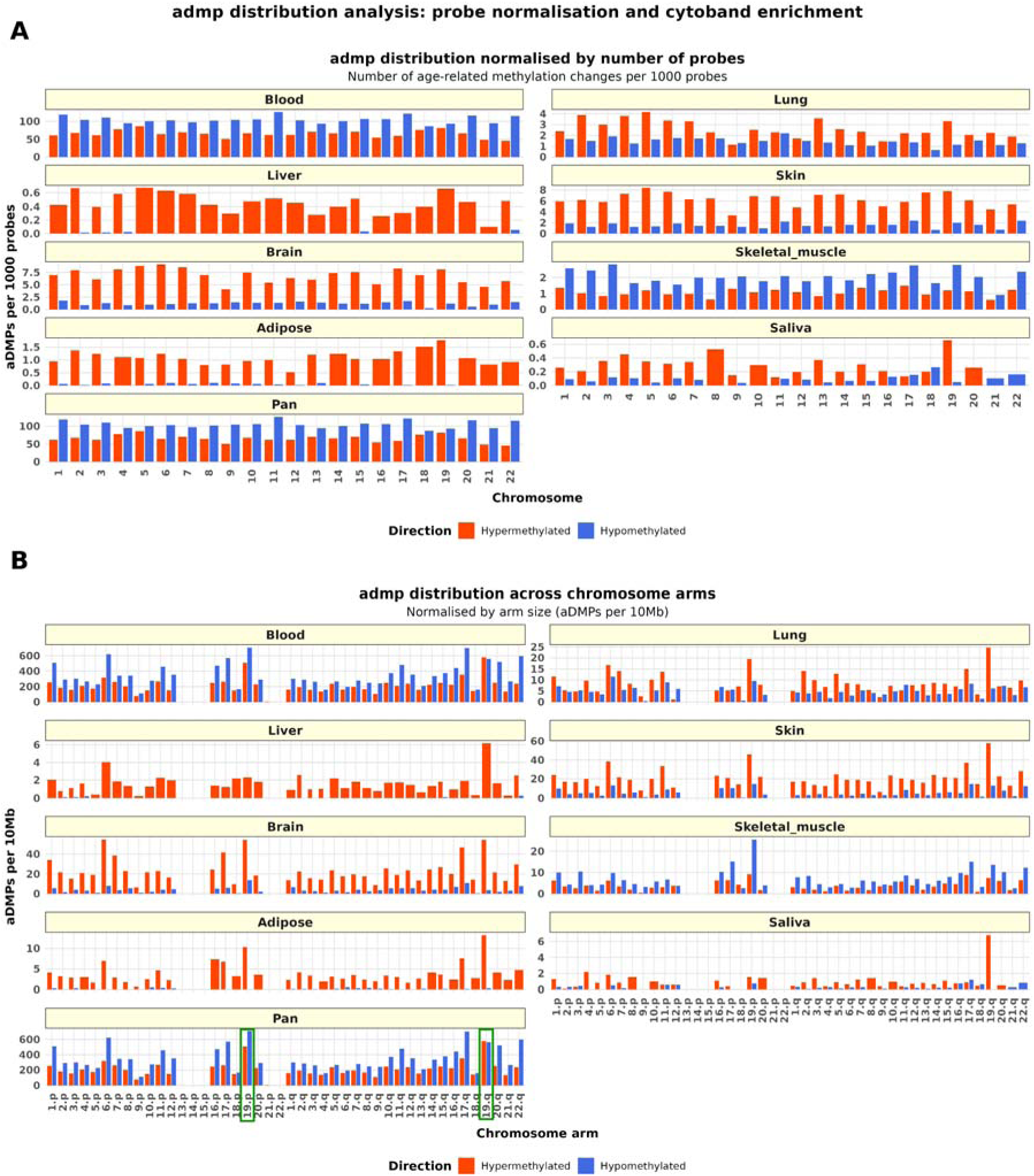
Chromosomal distribution of age-associated differentially methylated positions (aDMPs). (A) Number of hypermethylated and hypomethylated aDMPs per 1,000 probes across chromosomes, normalised for probe density. (B) Distribution of aDMPs across chromosome arms, normalised by arm length (aDMPs per 10 Mb). Chromosome 19 shows (highlighted in green box) consistent enrichment across multiple tissues and pan-tissue analyses.

**Supplementary Figure S7.**
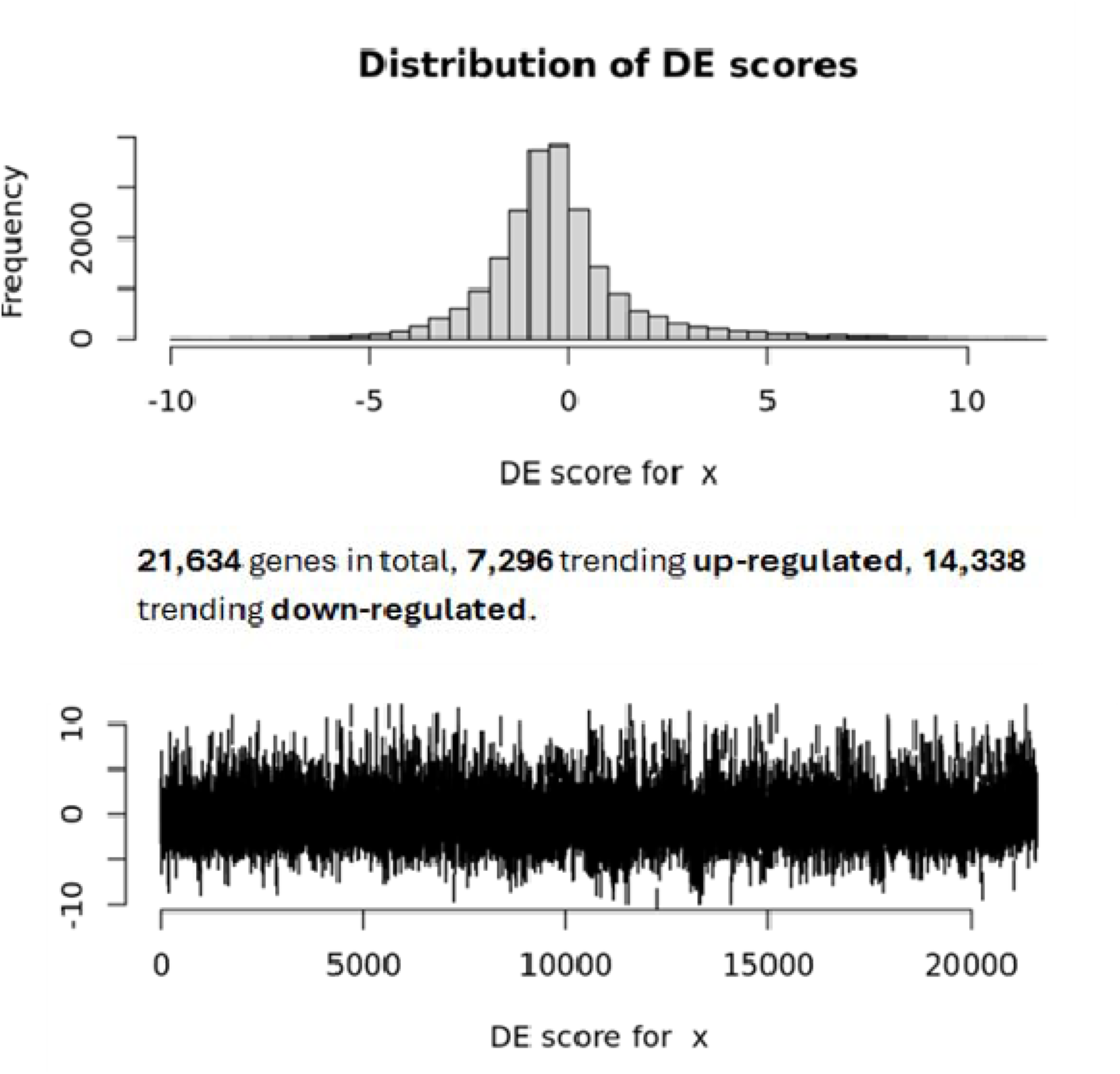
Gene-level directionality of age-associated methylation changes. Histogram and genome-wide distribution of differential expression (DE) scores derived from age-associated CpGs mapped to genes. Although fewer genes show hypermethylation than hypomethylation overall, hypermethylated genes cluster within specific pathways.

**Supplementary Figure S8.**
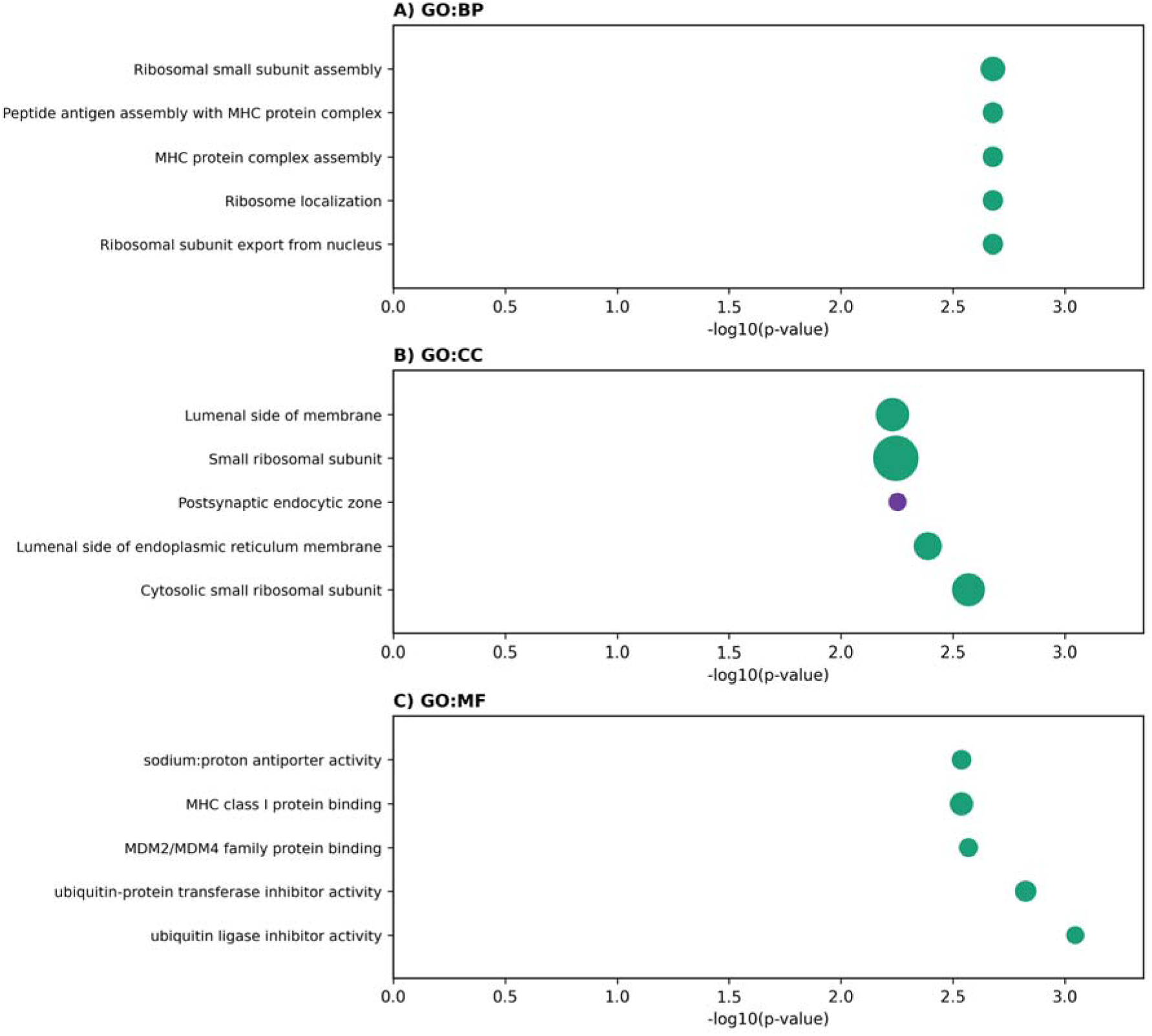
Top-five pathways ranked by −log[][](FDR) associated with age-related methylation changes in DMRs across pan-tissue analyses (FDR < 0.05). Pathways are shown for (A) Gene Ontology (GO) Biological Process (BP), (B) GO Cellular Component (CC), and (C) GO Molecular Function (MF). Only gene sets with a minimum size of n = 10 were included. Dot size reflects gene set size (range: N = 11–68). Colour indicates direction of enrichment (green = hypomethylation, purple = hypermethylation).

**Supplementary Figure S9.**
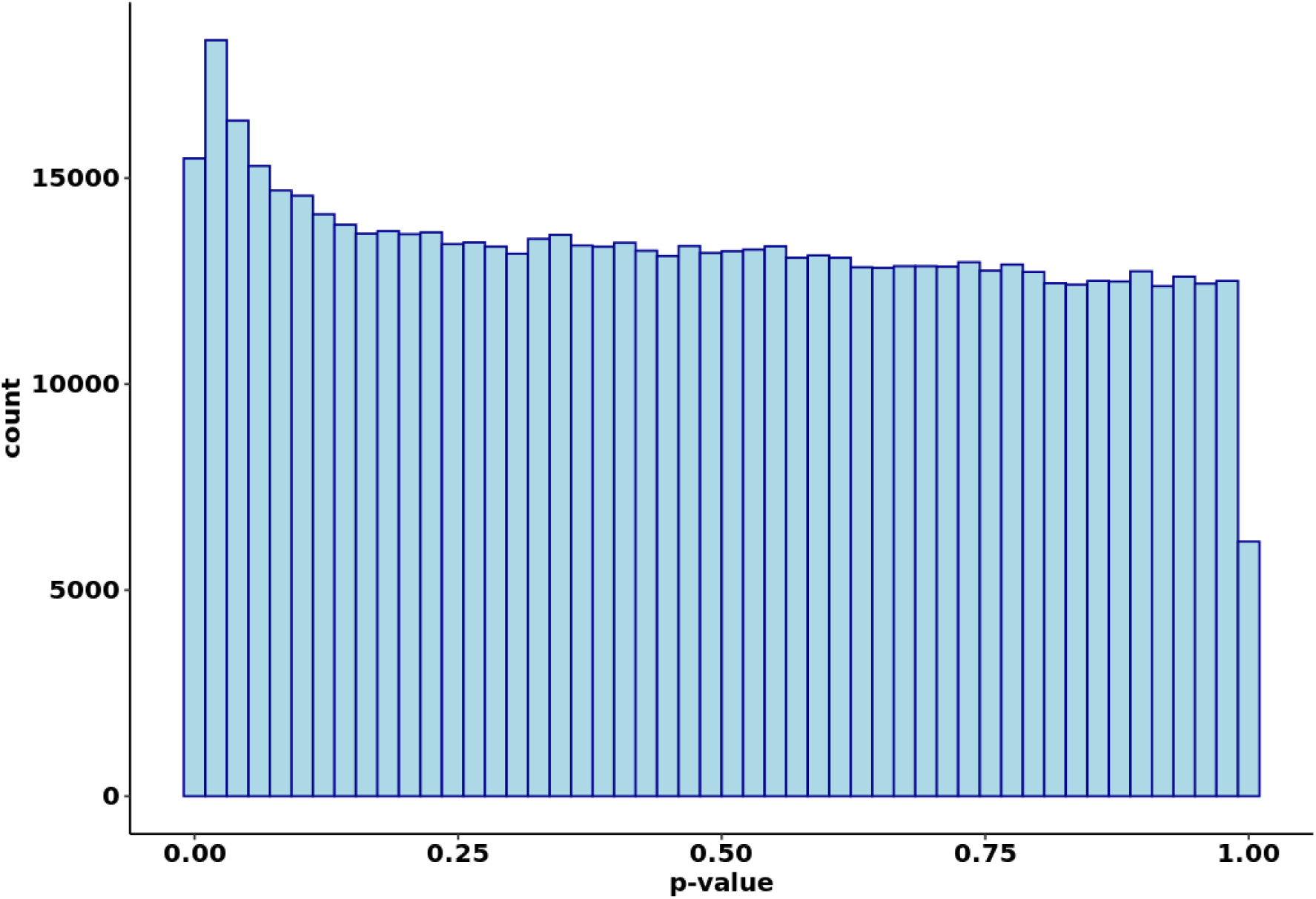
Distribution of raw p-values for sex-moderated age-associated DMPs. Histogram of unadjusted p-values from the meta-regression assessing sex moderation of the DNAm ∼ age relationship across pan-tissue datasets. Only CpGs with FDR < 0.05 were considered significant in downstream analyses.

**Supplementary Figure S10.**
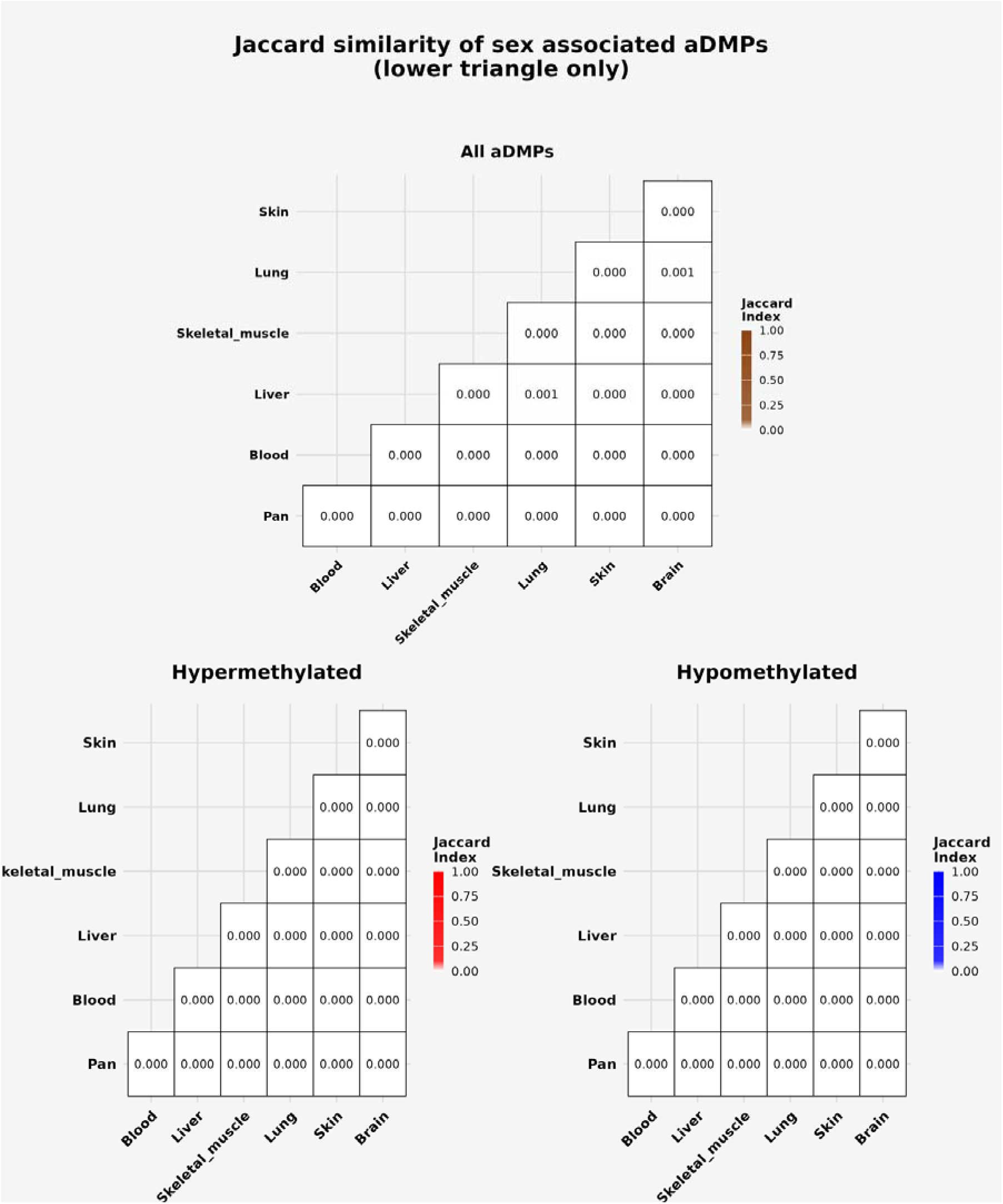
Jaccard similarity of sex-moderated aDMPs across tissues. Lower-triangle heatmaps showing pairwise Jaccard similarity indices for sex-moderated aDMPs across tissues. Top: all sex-moderated aDMPs. Bottom left: hypermethylated sites only. Bottom right: hypomethylated sites only. All indices equal zero, indicating complete tissue specificity of sex-moderated methylation changes.

**Supplementary Figure S11.**
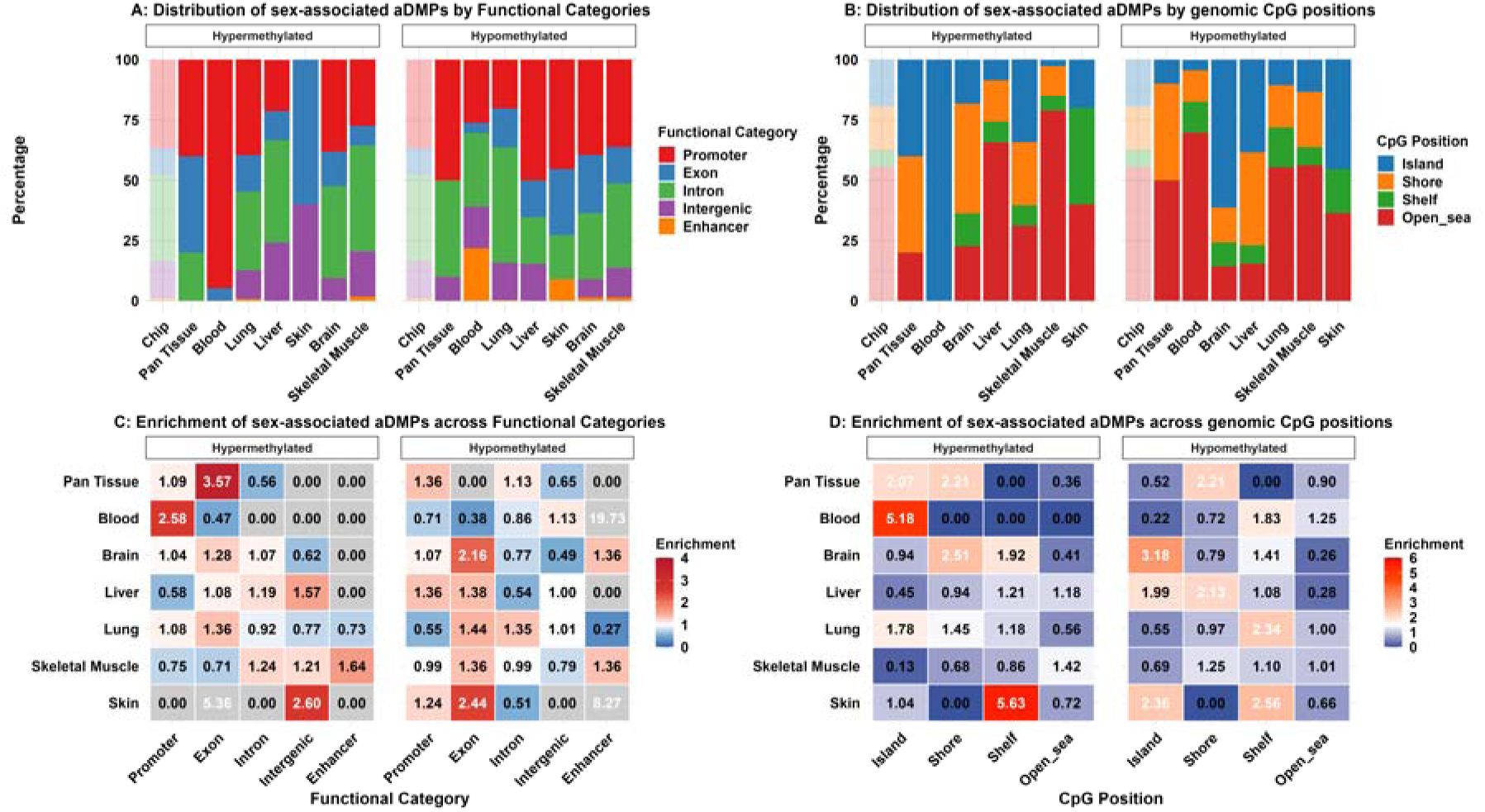
Functional genomic annotation of sex-moderated aDMPs. Distribution and enrichment of sex-moderated age-associated DMPs across CpG context (island, shore, shelf, open sea) and functional genomic regions (promoter, exon, intron, intergenic, enhancer), normalised to array probe distribution.

**Supplementary Figure S12.**
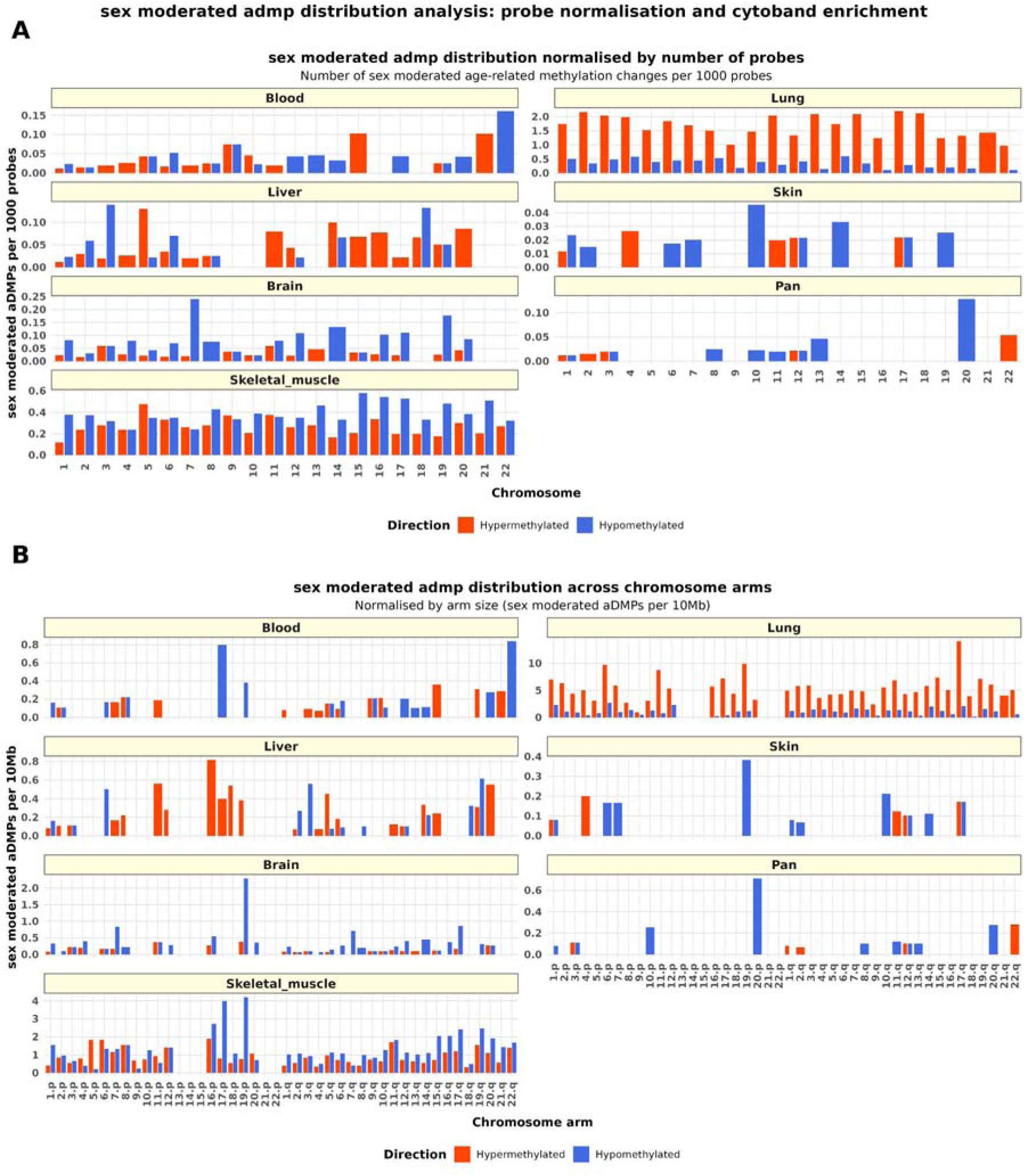
Chromosomal distribution of sex-moderated aDMPs. (A) Number of sex-moderated aDMPs per 1,000 probes across chromosomes. (B) Distribution of sex-moderated aDMPs across chromosome arms, normalised by arm size (per 10 Mb). The sparse distribution reflects the limited number of sex-moderated CpGs identified at FDR < 0.05.

## Supplementary Tables

**Supplementary Table 1.**
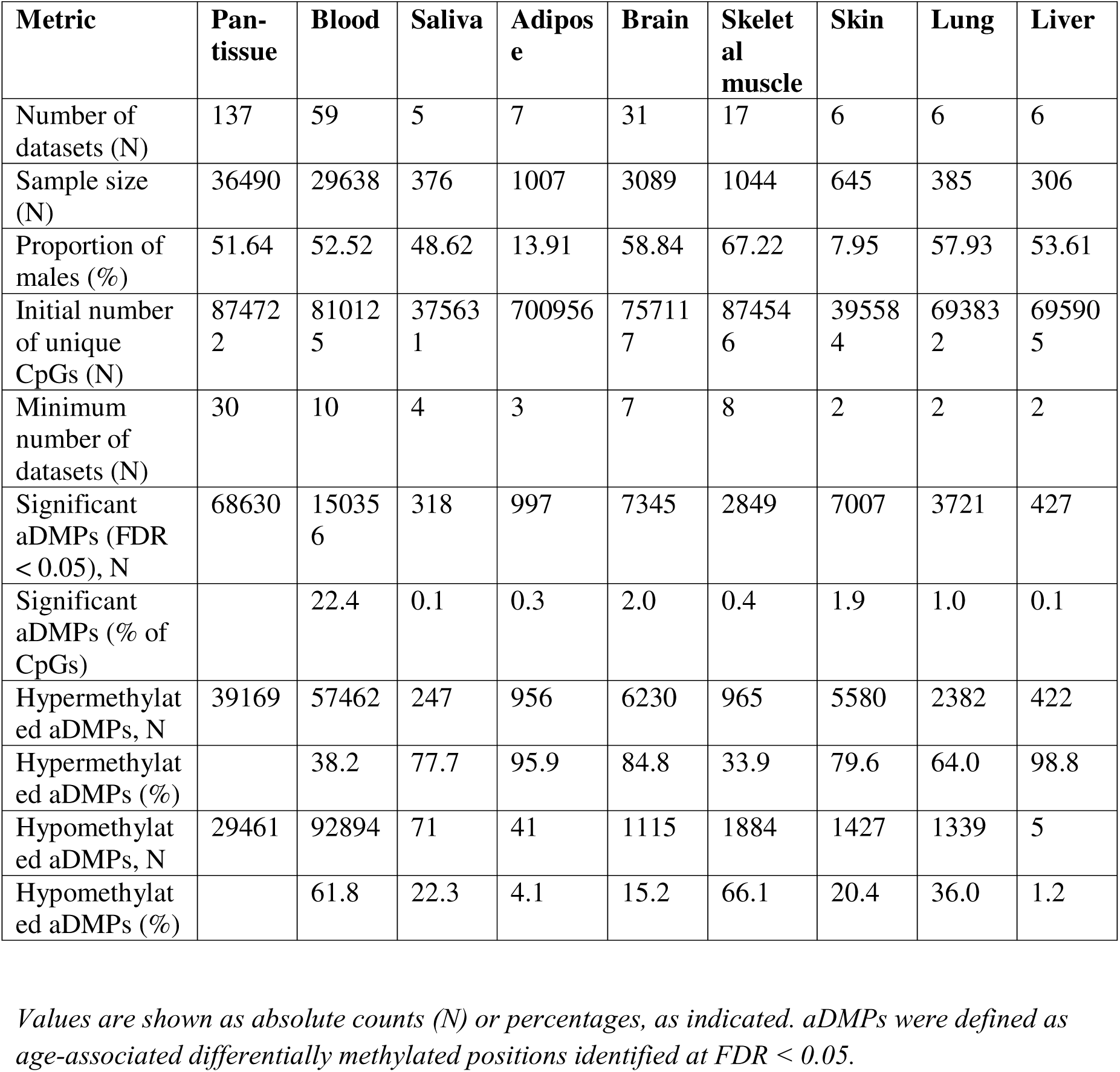
Summary characteristics of individual tissue meta-analyses.

**Supplementary Table 2a.** Distribution and enrichment of aDMPs across functional categories by tissue type and methylation direction at FDR<0.05.

**Background distribution of array probes**
| Category | Promoter (%) | Exon (%) | Intron (%) | Intergenic (%) | Enhancer (%) |
| --- | --- | --- | --- | --- | --- |
| Array probes | 36.7 | 11.2 | 35.5 | 15.4 | 1.1 |

**(A) Hypermethylated**
| Tissue type | Promoter % (E) | Exon % (E) | Intron % (E) | Intergenic % (E) | Enhancer % (E) |
| --- | --- | --- | --- | --- | --- |
| Blood | 48.8% (1.33) | 13.5% (1.21) | 25.1% (0.71) | 11.5% (0.75) | 1.2% (1.09) |
| Lung | 40.5% (1.10) | 19.5% (1.74) | 22.6% (0.64) | 16.3% (1.06) | 1.1% (1.00) |
| Liver | 50.6% (1.38) | 17.4% (1.55) | 21.1% (0.59) | 9.9% (0.64) | 1.0% (0.91) |
| Skin | 48.2% (1.31) | 16.5% (1.47) | 21.5% (0.61) | 12.4% (0.81) | 1.4% (1.27) |
| Brain | 50.6% (1.38) | 16.4% (1.46) | 20.1% (0.57) | 11.9% (0.77) | 1.0% (0.91) |
| Skeletal muscle | 31.5% (0.86) | 11.5% (1.03) | 44.8% (1.26) | 10.9% (0.71) | 1.3% (1.18) |

**(B) Hypomethylated**
| Tissue type | Promoter % (E) | Exon % (E) | Intron % (E) | Intergenic % (E) | Enhancer % (E) |
| --- | --- | --- | --- | --- | --- |
| Blood | 26.9% (0.73) | 11.7% (1.04) | 40.2% (1.13) | 19.6% (1.27) | 1.6% (1.45) |
| Lung | 23.7% (0.65) | 13.1% (1.17) | 43.9% (1.24) | 18.3% (1.19) | 1.0% (0.91) |
| Liver | 20.0% (0.54) | 20.0% (1.79) | 20.0% (0.56) | 40.0% (2.60) | 0.0% (0.00) |
| Skin | 36.0% (0.98) | 14.1% (1.26) | 34.5% (0.97) | 14.1% (0.92) | 1.2% (1.09) |
| Brain | 33.4% (0.91) | 14.5% (1.29) | 40.6% (1.14) | 10.0% (0.65) | 1.4% (1.27) |
| Skeletal muscle | 29.8% (0.81) | 20.7% (1.85) | 40.0% (1.13) | 8.9% (0.58) | 0.6% (0.55) |

*Percentages indicate the proportion of aDMPs within each functional category.*
*Enrichment values (E) represent observed-to-expected ratios relative to the background distribution of array probes. aDMPs were defined at FDR < 0.05.*

**Supplementary Table 2b.** Distribution and enrichment of aDMPs across genomic location by tissue type and methylation direction at FDR<0.05.

**Background distribution of array probes**
|  | Island | Shore | Shelf | Open sea |
| --- | --- | --- | --- | --- |
| Array probes | 19.3% | 18.1% | 7.1% | 55.5% |

(A) Hypermethylated
| Tissue type | Island % (E) | Shore % (E) | Shelf % (E) | Open sea % (E) |
| --- | --- | --- | --- | --- |
| Blood | 100.0 (5.18) | 0.0 (0.00) | 0.0 (0.00) | 0.0 (0.00) |
| Lung | 34.3 (1.78) | 26.2 (1.45) | 8.4 (1.18) | 31.1 (0.56) |
| Liver | 8.7 (0.45) | 17.0 (0.94) | 8.6 (1.21) | 65.5 (1.18) |
| Skin | 20.0 (1.04) | 0.0 (0.00) | 40.0 (5.63) | 40.0 (0.72) |
| Brain | 18.2 (0.94) | 45.5 (2.51) | 13.6 (1.92) | 22.7 (0.41) |
| Skeletal muscle | 2.5 (0.13) | 12.3 (0.68) | 6.1 (0.86) | 78.9 (1.42) |

(B) Hypomethylated
| Tissue type | Island % (E) | Shore % (E) | Shelf % (E) | Open sea % (E) |
| --- | --- | --- | --- | --- |
| Blood | 4.2 (0.22) | 13.0 (0.72) | 13.0 (1.83) | 69.4 (1.25) |
| Lung | 10.6 (0.55) | 17.6 (0.97) | 16.6 (2.34) | 55.5 (1.00) |
| Liver | 38.4 (1.99) | 38.5 (2.13) | 7.7 (1.08) | 15.5 (0.28) |
| Skin | 45.5 (2.36) | 0.0 (0.00) | 18.2 (2.56) | 36.6 (0.66) |
| Brain | 61.4 (3.18) | 14.3 (0.79) | 10.0 (1.41) | 14.4 (0.26) |
| Skeletal muscle | 13.3 (0.69) | 22.6 (1.25) | 7.8 (1.10) | 56.1 (1.01) |

Percentages indicate the proportion of aDMPs within each CpG position category. Enrichment values (E) represent observed-to-expected ratios relative to the background distribution of array probes. aDMPs were defined at FDR < 0.05.

**Supplementary Table 3a.** Details of the top five statistically significant pathways associated with age-related methylation changes in DMPs ranked by smallest enrichment score (S score). Minimum gene set of n = 10.

|  | Pathway | Set size (N>10) | S. distance /Enrichment score | FDR ANOVA |
| --- | --- | --- | --- | --- |
| Reactome | Specification of the neural plate border. | 16 | 0.828 | 9.53 x 10<sup>-7</sup> |
|  | Formation of the nephric duct | 17 | 0.7710 | 2.45x 10<sup>-6</sup> |
|  | Formation of the ureteric bud | 21 | 0.7361 | 4.30x 10<sup>-7</sup> |
|  | FGFRL1 modulation of FGFR1 signaling | 13 | 0.7294 | 2.08x 10<sup>-4</sup> |
|  | Kidney development | 45 | 0.7186 | 1.30e- x 10<sup>-14</sup> |
| KEGG: | Lectin pathway of the coagulation cascade (fibrinogen to fibrin) | 10 | -0.659 | 6.89 × 10<sup>-3</sup> |
|  | Wnt signalling modulation via LGR-RSPO | 19 | 0.640 | 1.04 × 10<sup>-4</sup> |
|  | MLL-ENL fusion-driven transcriptional activation | 12 | 0.612 | 5.74 × 10<sup>-3</sup> |
| | IGH mmset fusion to transcriptional activation | 16 | 0.60 | $8.01 \times 10^{-4}$ |
| | WNT signaling modulation wnt acylation | 18 | 0.59 | $4.06 \times 10^{-4}$ |
| GO:BP | Pancreatic $\alpha$ -cell differentiation | 10 | 0.95 | $3.98 \times 10^{-6}$ |
| | Nephric duct morphogenesis | 10 | 0.89 | $1.70 \times 10^{-5}$ |
| | Mesonephric tubule formation | 10 | 0.86 | $3.51 \times 10^{-5}$ |
| | Nephron tubule formation | 20 | 0.86 | $1.47 \times 10^{-9}$ |
| | Spinal cord association neuron differentiation | 13 | 0.79 | $1.28 \times 10^{-5}$ |
| GO:CC | Platelet dense granule lumen | 14 | -0.63 | $5.81 \times 10^{-4}$ |
| | Muscle myosin complex | 15 | -0.56 | $1.65 \times 10^{-3}$ |
| | Sarcoplasmic reticulum lumen | 10 | -0.56 | $1.80 \times 10^{-2}$ |
| | Chromosome centromeric core domain | 16 | 0.55 | $1.65 \times 10^{-3}$ |
| | Voltage gated sodium channel complex | 14 | -0.48 | $1.70 \times 10^{-2}$ |
| GO:MF | HMG box domain binding | 12 | 0.810 | $7.57 \times 10^{-5}$ |
| | Adrenergic receptor activity | 10 | 0.77 | $7.30 \times 10^{-4}$ |
| | Catecholamine binding | 11 | 0.75 | $4.60 \times 10^{-4}$ |
| | Odorant binding | 82 | -0.69 | $5.57 \times 10^{-25}$ |
| | Co receptor binding | 13 | 0.63 | $1.78 \times 10^{-3}$ |

**Supplementary Table 3b.** Details of the top five statistically significant pathways associated with age-related methylation changes in DMRs ranked by smallest enrichment score (S score). Minimum gene set of n = 10.

|  | Pathway | Set size (N>10) | Direction | Non-adjusted p-value |
| --- | --- | --- | --- | --- |
| GO:BP | Ribosomal subunit export from nucleus | 14 | Hypo | 0.0021 |
|  | Ribosome localization | 14 | Hypo | 0.0021 |
|  | MHC protein complex assembly | 14 | Hypo | 0.0021 |
|  | Peptide antigen assembly with MHC protein complex | 14 | Hypo | 0.0021 |
|  | Ribosomal small subunit assembly | 20 | Hypo | 0.0021 |
| GO:CC | Cytosolic small ribosomal subunit | 36 | Hypo | 0.0027 |
|  | Lumenal side of endoplasmic reticulum membrane | 26 | Hypo | 0.0041 |
|  | Postsynaptic endocytic zone | 11 | Hyper | 0.0056 |
|  | Small ribosomal subunit | 68 | Hypo | 0.0057 |
|  | Lumenal side of membrane | 37 | Hypo | 0.0059 |
| GO:MF | ubiquitin ligase inhibitor activity | 11 | Hypo | 0.0009 |
|  | ubiquitin-protein transferase inhibitor activity | 15 | Hypo | 0.0015 |
|  | MDM2/MDM4 family protein binding | 12 | Hypo | 0.0027 |
|  | MHC class I protein binding | 18 | Hypo | 0.0029 |
|  | sodium:proton antiporter activity | 13 | Hypo | 0.0029 |

**Supplementary Table 4a.** Distribution and enrichment of sex-moderated aDMPs across functional categories by tissue type and methylation direction at FDR<0.05.

**Background distribution of array probes**
| Category | Promoter (%) | Exon (%) | Intron (%) | Intergenic (%) | Enhancer (%) |
| --- | --- | --- | --- | --- | --- |
| Array probes | 36.7 | 11.2 | 35.5 | 15.4 | 1.1 |

**(A) Hypermethylated**
| Tissue type | Promoter % (E) | Exon % (E) | Intron % (E) | Intergenic % (E) | Enhancer % (E) |
| --- | --- | --- | --- | --- | --- |
| Blood | 94.70% (2.58) | 5.30% (0.47) | 0.00% (0.00) | 0.00% (0.00) | 0.00% (0.00) |
| Lung | 39.60% (1.08) | 15.20% (1.36) | 32.60% (0.92) | 11.90% (0.77) | 0.80% (0.73) |
| Liver | 21.20% (0.58) | 12.10% (1.08) | 42.40% (1.19) | 24.20% (1.57) | 0.00% (0.00) |
| Skin | 0.00% (0.00) | 60.00% (5.36) | 0.00% (0.00) | 40.00% (2.60) | 0.00% (0.00) |
| Brain | 38.10% (1.04) | 14.30% (1.28) | 38.10% (1.07) | 9.50% (0.62) | 0.00% (0.00) |
| Skeletal muscle | 27.60% (0.75) | 8.00% (0.71) | 44.00% (1.24) | 18.70% (1.21) | 1.80% (1.64) |

**(B) Hypomethylated**
| Tissue type | Promoter % (E) | Exon % (E) | Intron % (E) | Intergenic % (E) | Enhancer % (E) |
| --- | --- | --- | --- | --- | --- |
| Blood | 26.10% (0.71) | 4.30% (0.38) | 30.40% (0.86) | 17.40% (1.13) | 21.70% (19.73) |
| Lung | 20.20% (0.55) | 16.10% (1.44) | 47.90% (1.35) | 15.50% (1.01) | 0.30% (0.27) |
| Liver | 50.00% (1.36) | 15.40% (1.38) | 19.20% (0.54) | 15.40% (1.00) | 0.00% (0.00) |
| Skin | 45.50% (1.24) | 27.30% (2.44) | 18.20% (0.51) | 0.00% (0.00) | 9.10% (8.27) |
| Brain | 39.40% (1.07) | 24.20% (2.16) | 27.30% (0.77) | 7.60% (0.49) | 1.50% (1.36) |
| Skeletal muscle | 36.20% (0.99) | 15.20% (1.36) | 35.00% (0.99) | 12.20% (0.79) | 1.50% (1.36) |

*Percentages indicate the proportion of sex-moderated aDMPs within each functional category. Enrichment values (E) represent observed-to-expected ratios relative to the background distribution of array probes. aDMPs were defined at FDR < 0.05.*

**Supplementary Table 4b.** Distribution and enrichment of sex-moderated aDMPs across genomic location by tissue type and methylation direction at FDR<0.05.

**Background distribution of array probes**
|  | **Island** | **Shore** | **Shelf** | **Open sea** |
| --- | --- | --- | --- | --- |
| Array probes | 19.3% | 18.1% | 7.1% | 55.5% |

**(A) Hypermethylated**
| **Tissue type** | **Island count (%)** | **Shore count (%)** | **Shelf count (%)** | **Open sea count (%)** |
| --- | --- | --- | --- | --- |
| Blood | 31073 (54.2%) | 14445 (25.2%) | 1737 (3.0%) | 10127 (17.6%) |
| Lung | 1477 (62.1%) | 688 (28.9%) | 57 (2.4%) | 157 (6.6%) |
| Liver | 228 (54.0%) | 148 (35.1%) | 12 (2.8%) | 34 (8.1%) |
| Skin | 3475 (62.4%) | 1337 (24.0%) | 129 (2.3%) | 631 (11.3%) |
| Brain | 3720 (59.8%) | 1687 (27.1%) | 166 (2.7%) | 650 (10.4%) |
| Skeletal muscle | 139 (14.4%) | 180 (18.7%) | 94 (9.7%) | 552 (57.2%) |

**(B) Hypomethylated**
| **Tissue type** | **Island count (%)** | **Shore count (%)** | **Shelf count (%)** | **Open sea count (%)** |
| --- | --- | --- | --- | --- |
| Blood | 3674 (4.0%) | 21623 (23.3%) | 10062 (10.9%) | 57353 (61.9%) |
| Lung | 59 (4.4%) | 373 (27.9%) | 199 (14.9%) | 707 (52.8%) |
| Liver | 0 (0.0%) | 2 (40.0%) | 0 (0.0%) | 3 (60.0%) |
| Skin | 63 (4.4%) | 624 (43.8%) | 203 (14.2%) | 535 (37.5%) |
| Brain | 86 (7.7%) | 383 (34.4%) | 156 (14.0%) | 489 (43.9%) |
| Skeletal muscle | 115 (6.1%) | 524 (27.8%) | 234 (12.4%) | 1011 (53.7%) |

*Percentages indicate the proportion of sex-moderated aDMPs within each CpG position category. Enrichment values (E) represent observed-to-expected ratios relative to the background distribution of array probes. aDMPs were defined at FDR < 0.05.*

**Supplementary Table 5.**
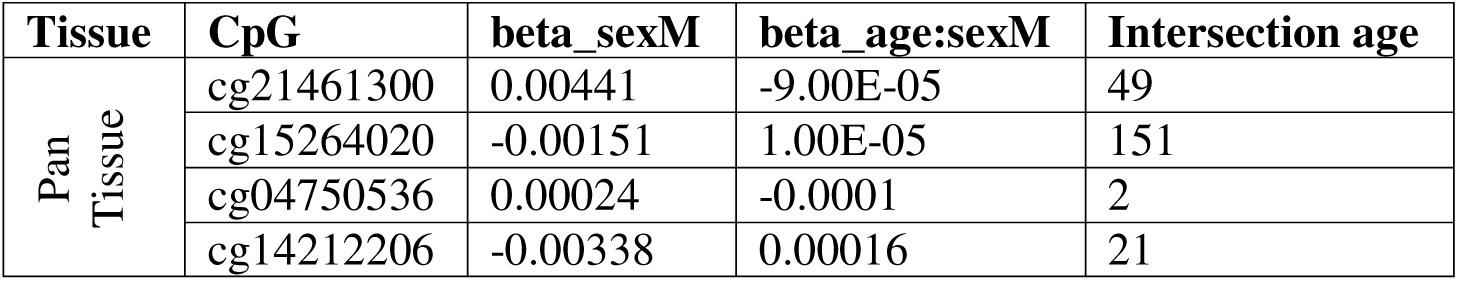
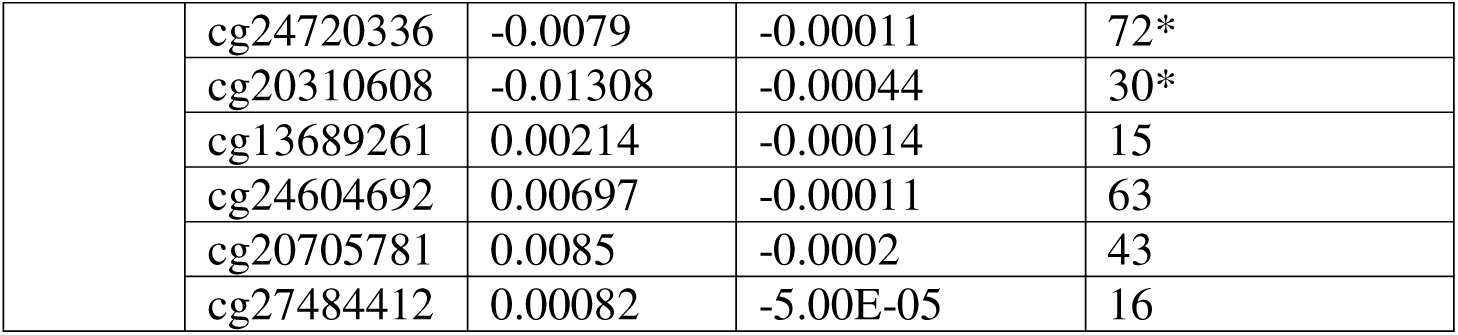
Summary characteristics of interactive term modelling of the top 10 sex-associated aDMPs.

